# A Single Cell Atlas of the Newt Iris During Lens Regeneration

**DOI:** 10.64898/2025.12.06.692619

**Authors:** Olivia M. Williams, Kelsey E. Ahearn, Joseph L. Sevigny, Nicole Farber, Disha Hegde, Kenneth J. Lampel, Jenna Loporcaro, Leo Napoleon, Jacob Nipoti, Timothy Ralich, Brooklyn Wallace, W. Kelley Thomas, Konstantinos Sousounis

## Abstract

Iris pigmented epithelial (IPE) cells transdifferentiate to lens epithelial cells (LECs) during Wolffian lens regeneration in newts. Single cell RNA sequencing was used at multiple timepoints to further our understanding of this process and the cells involved in it. All major cell types present in and adjacent to the iris were identified including IPE cells, macrophages, non-pigmented ciliary epithelial cells, pigmented ciliary epithelial cells, and stroma-residing fibroblasts, endothelial cells, iridophores, and melanocytes. In the intact iris, IPE cell subpopulations were characterized by the expression of the dorsoventral genes TBX5 and VAX2, and newly identified markers LTBP2, CHRM3, and NTN1. During regeneration, IPE heterogeneity was correlated with functional states such as the cell cycle, migration, and lens vesicle formation. Pseudotime trajectory analysis revealed new insights into transcriptional and reprogramming factors during the IPE-to-LEC conversion and built a molecular and genetic blueprint of newt lens regeneration. Macrophages were identified as tissue-resident and underwent polarization from M1 early to M2 late during lens regeneration, an event that correlated temporally with the IPE-to-LEC reprogramming. Overall, this atlas provides data and analysis for iris cell types, IPE subpopulations, IPE cell states, gene expression changes as IPE cells reprogram to LECs, macrophage identity and function, and cell-to-cell interactions during newt lens regeneration.

**Highlights:**

- Cell atlas identifying cells in the newt iris at multiple timepoints during lens regeneration
- Intact iris contains multiple iris pigmented epithelial (IPE) cell subpopulations
- Identification of IPE functional states during regeneration
- Cellular trajectory analysis revealed a molecular and genetic blueprint of IPE-to-lens epithelial cell reprogramming
- Identification of cell-to-cell interactions between IPE cells and other cell types
- Macrophages interacting with IPE cells are tissue-resident and polarize from M1 to M2 subtypes during lens regeneration

**Graphical Abstract:** 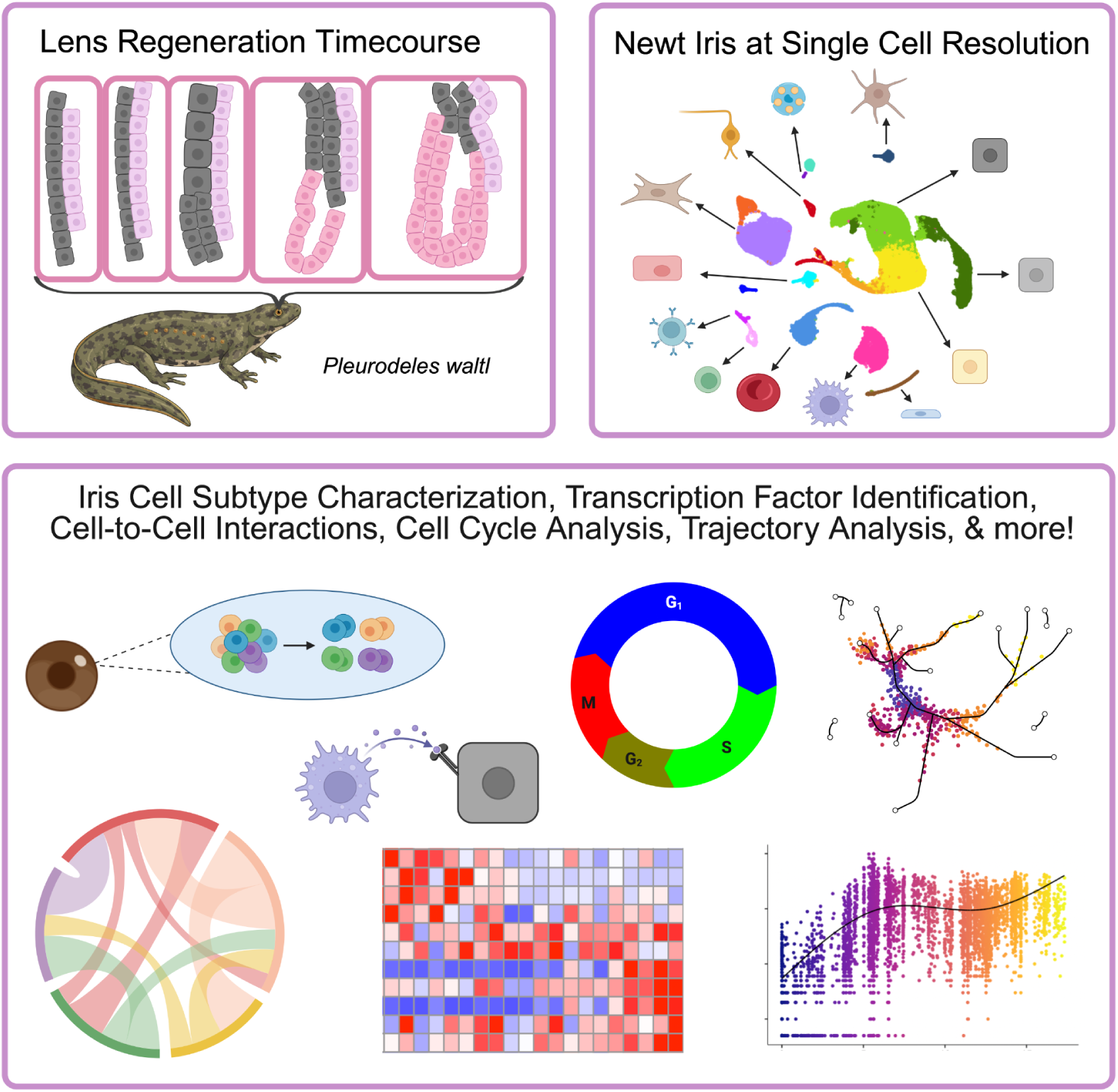

## Introduction

Newts can regenerate their eye lens following its complete removal throughout life^1–3^. To accomplish this, pigmented epithelial cells residing in the dorsal iris transdifferentiate to lens epithelial cells (LECs) and regenerate the missing lens. Termed Wolffian lens regeneration, this process is unique in sourcing the neuroectodermally-derived iris pigmented epithelial (IPE) cells to regenerate the originally surface ectoderm-derived lens. This deviation from the normal developmental fate trajectories is not typical in regeneration models, like salamander limb and mouse digit tip regeneration, which typically rely on their lineage to restore tissues^4,5^. Even frog tadpoles that regenerate their lens before metamorphosis rely on their corneal epithelium which is derived from the surface ectoderm like the lens^6^. Given these, studying the transdifferentiation process during newt lens regeneration may unravel interesting and novel biological mechanisms.

Following the complete removal of the newt eye lens, IPE cells enlarge, enter the cell cycle, depigment, and form a vesicle comprised of LECs in the anterior and fibers in the posterior. This vesicle grows over time to replace the lost lens^3^. Microarrays, bulk RNA sequencing, and mass spectrometry have been used in the past to study how gene expression changes during the IPE-to-LEC reprogramming process. These approaches have increased our understanding of newt lens regeneration by revealing potential roles for the T-box transcriptional factor TBX5 in the dorsal iris, the Ventral anterior homeobox 2 (VAX2) in the ventral iris, extracellular matrix proteins like fibrillin, signaling pathways like Netrin and Ephrin, and molecular events like DNA damage and immune response during the process among others ^7–9^. However, these studies had limitations including the use of bulk tissue and the relatively few timepoints sampled typically concentrated at the early stages of regeneration. At the cellular level, it is well documented that IPE cells are responsible for reprogramming to LECs, however, they are not well characterized at the molecular level, and it is unclear whether different subpopulations of IPE cells are present in the newt iris. Macrophages is another cell population with known roles during lens regeneration. Shortly after lens removal, macrophages significantly increase in number near the dorsal iris and scavenge the secreted pigment granules from IPE cells that undergo dedifferentiation^10,11^. They are also thought to engage in the immune response and cell signaling by secreting ligands^12^. The exact role and cellular identity of these macrophages are still unclear.

To fill these gaps, we performed a single cell RNA sequencing (scRNAseq) experiment to explore in detail the lens regeneration process in the genetically tractable newt *Pleurodeles waltl*. We used 10 different timepoints spanning from intact iris to 16 days post lens removal (dpl) time that the lens vesicle has formed in juvenile animals. We sequenced the samples very deeply to extract as much transcriptomic information as possible so this dataset could serve as a useful resource for the field. We took advantage of the recently sequenced and annotated genome of *Pleurodeles waltl* to base our downstream *in silico* analysis^13^. Our study provides new data for (a) IPE cell heterogeneity in the intact iris, (b) gene expression patterns, cellular functions, and different cell states of IPE cells following lens removal, (c) cellular trajectories and gene regulation that governs fate transitions during IPE-to-LEC reprogramming, and (d) functions of other cells including new insights into macrophages during lens regeneration.

## Results

### Cellular heterogeneity of the newt iris

Lens regeneration undergoes distinct histologically-defined stages starting with the thickening of the iris (Sato stages 1 and 2, Reyer stage I; latent period) and followed by IPE depigmentation and lens vesicle formation (Sato stages 3-6, Reyer stage II; initial period). These events mark the end of the IPE-to-LEC reprogramming and the initiation of fiber differentiation by LECs and the subsequent growth of the lens (Sato stages 7-13, Reyer stages III and IV)^3^. To make this atlas, ten different timepoints encompassing Reyer stages I and II were surveyed to achieve high temporal resolution of the IPE cell transdifferentiation process (Figure 1A and 1B). Lens removals were performed in same-clutch sibling *Pleurodeles waltl* newts followed by eye and iris extractions. Samples were enriched for the lens regeneration-competent dorsal iris. Tissues were dissociated into single cells, fixed following the Parse Biosciences protocol and stored until all samples were collected. Single cell barcoding and library preparation were performed using the Parse Biosciences platform followed by sequencing and alignment to the *Pleurodeles waltl* genome^13^ (Figure 1C). 86,460 cells were identified and sequenced on average at 101,958 reads per cell with a median of 3,542 genes recovered and 72.7% gene discovery saturation. Following filtering through quality control parameters including cell size distribution, number of genes, and doublet removal, 80,425 cells remained for downstream analysis (Table S1). Uniform Manifold Approximation and Projection (UMAP) analysis was used to group and identify the cellular heterogeneity of the newt iris (Figure 1D and Figure S1A). Since newt IPE cells are unique in their ability to reprogram to LECs, and the whole dataset included previously uncharacterized IPE derivatives and cell states assumed during regeneration, it was expected that IPE cells may present novel expression profiles at different timepoints. Relying on well-documented anterior eye segment cell type marker genes from other vertebrates we identified 13 cell types; L1CAM+ Schwann cells, CD34+ endothelial cells, CSF1R+ macrophages, CD78B+ B cells, CYTIP+ T cells, HBB+ erythrocytes, LTK+ iridophores, PAX3+ melanocytes, PLEK+ platelets, ENPP6+ iris sphincter cells, FBN2+ non-pigmented ciliary epithelial (NPCE) cells, GLIS1+ fibroblasts, and PAPPA+ pigmented ciliary epithelial (PCE) cells^14–18^ (Figure 1E and Figure S1B). UNC80+ and GLDN+ retina cells and KRT12+ cornea cells were also captured due to tissue cross-contamination during iris extraction (Figure 1E and Figure S1B). IPE cells were initially defined by the expression of the pigment marker OCA2^19^ and the low expression of the PCE marker PAPPA and the NPCE marker FBN2. Cells with this profile highly expressed the gene LTBP2 consistently throughout the timepoints relative to all other cell types and was used as an IPE marker (Figure 1E, Table S2, and Figure S1B). IPE, PCE, and NPCE cells were positive for the eye neuroepithelial transcriptional factor PAX6 (Figure S1C). All expected anterior eye segment cell types appeared in all timepoints sampled but their distribution dynamically changed during lens regeneration (Figure 1F). IPE cells and macrophages were increasingly more abundant overtime following lens removal compared to other cell types as indicated by Generalized Linear Models analysis (GLM; Figure S1D). These data were in line with histological observations shown that IPE cells proliferate and macrophages increase their presence in the dorsal iris^11^ following lens removal further indicating that our experiment has faithfully captured the cellular heterogeneity of the newt iris during lens regeneration.

**Figure 1:**
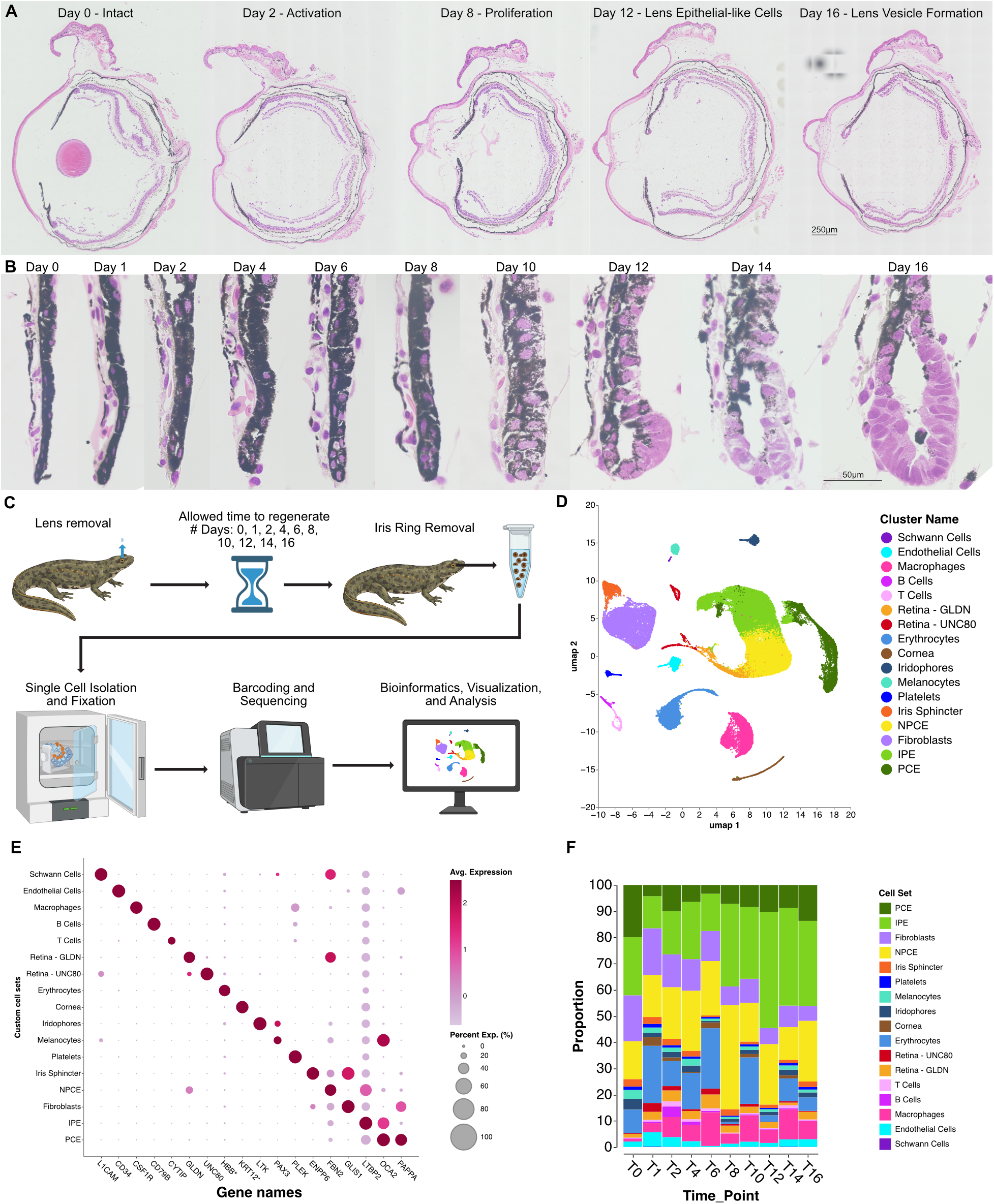
The cellular heterogeneity of the newt iris. A) Eye sections stained with H&E at 0-, 2-, 8-, 12-, and 16-days post lens removal. Scale bar: 250μm. B) High magnification of the dorsal iris across all timepoints. Scale bar: 50μm. C) Cartoon schematic of experimental design generated with BioRender. D) UMAP of all cells colored for custom cell sets. E) Dot plot of marker genes for different cell types identified. F) Frequency plot of cell types across timepoints.

Overall, our scRNAseq experiment has captured all predicted cell types present in the iris, iris stroma, and adjacent tissues, and provided molecular markers for the identification of the newt IPE cells.

### Distinct molecular stages during lens regeneration

First, general trends of gene expression changes were analyzed using Pearson correlation across all timepoints. IPE, NPCE, PCE, fibroblasts, and macrophages responded to lens removal relatively more than other cell types which largely appeared unaffected (Figure S2A). IPE cells, the cellular source of lens regeneration, had the most changes in gene expression followed by NPCE and macrophages (Figure 2A and Figure S2A). This analysis also revealed three distinct stages, intact, early (1-8dpl), and late (10-16dpl) (Figure 2A). The early and late regeneration stages, defined at the molecular level by the scRNAseq, were consistent with the histological Reyer stages I and II, respectively^3^, assessed by sibling- and timepoint-matched eyes (Figure 1A and 1B). Lens regeneration-competent dorsal IPE cells were further identified based on the lack of the ventral marker gene VAX2. The majority of these cells were enriched for the dorsal marker TBX5 (Figure S2B). Both dorsal and ventral IPE cells significantly changed their gene expression within one day following lens removal (Figure 2B). However, principal component analysis (PCA) showed that dorsal IPE cells changed relatively more than ventral. Specifically, 14dpl ventral IPE cells had a comparable response as 4dpl dorsal IPE cells (Figure 2C). These data corroborated previous reports that ventral IPE cells have delayed progression towards a LEC fate^20,21^. Gene Ontology (GO) enrichment analysis for genes expressed in dorsal and ventral IPE during lens regeneration, revealed stage-specific functions (Table S4). The intact iris expressed genes involved in adherens junctions, signal transduction, and postsynaptic membrane (Figure 2D; light blue and green bars). Adherens junctions are a major component of various epithelia and functions to ensure shape and overall structure integrity with our data indicating similar roles in the iris^22^. Signal transduction and postsynaptic membrane categories included genes related to neuronal functions, receptors and ion transporters, which overall indicated the developmental neuroepithelial origin of the IPE cells. Early following lens removal, IPE cells expressed genes indicating increased translation, ribosomal assembly, and rRNA processing (Figure 2D; blue and green bars). These functions reflected the overall size increase for IPE cells at this stage (Figure 1B). IPE cells also upregulated genes related to focal adhesions which indicated a switch from the static adherens junctions to a pro-migratory state^23^. Dorsal IPE cells were enriched for genes associated with DNA replication, fibrillar center, and the mitochondrion (Figure 2D; blue bars). This suggested that dorsal IPE cells entered the cell cycle, produced proteins, and generated energy at greater levels than ventral. These functions could be associated with the overall faster gene expression response for dorsal compared to ventral IPE cells. These data were also corroborated with cell cycle phase analysis which indicated that dorsal IPE cells entered the S phase faster and at greater numbers (Figure 2E and Figure S2C). For instance, following lens removal, about 33% of dorsal IPE cells entered the S phase as early as Day 2 and peaked at Day 12 at 60% compared to 12% and 50% for ventral IPE cells, respectively. During the late stage, IPE cells expressed axon guidance genes with dorsal IPE cells being enriched for the axoneme (Figure 2D). This indicated that while IPE cells are not known to function as neurons or have axons, they utilized cellular machinery related to their neuroepithelial origin for migration towards or away from the developing lens vesicle. On the other hand, ventral IPE cells were enriched for Ephrin signaling, a mechanism known for eye segmentation^24^ (Figure 2D; dark green bar). Dorsal IPE cells were also enriched for genes associated with G couple-protein activity and collagen-containing extracellular matrix (Figure 2D; dark blue bars). These functions reflected the active intracellular signaling required for IPE-to-LEC reprogramming and the generation of the collagen-rich lens capsule found at this stage. To further understand how gene expression may be regulated during the different molecular stages, we performed transcriptional factor prediction analysis using Enrichr for the genes overrepresented in dorsal IPE cells^25^. The intact stage was found to be mainly regulated by the nuclear and hormone receptors RXRA, ESRRA, NR0B1, NR1I2, and NCOA1 (Figure 2F; Intact). During the early stage, gene expression was regulated by the chromatin remodeling factors SPI1 and EP300 and the cell differentiation factors BATF and LMO2, indicating the reprogramming of the IPE cell fate. JUN, a transcriptional factor activated downstream of FGF signaling a pathway known to be engaged during lens regeneration^20^, was also predicted to regulate genes at the early stage (Figure 2F; Early). The late stage was mainly regulated by developmental and cell differentiation genes including FOXP2, EMX2, KLF2, GFI1B, HOXD13, and MITF (Figure 2F; Late). These genes reflected the balance between IPE fate retainment or LEC fate switch commitment found at this stage.

**Figure 2:**
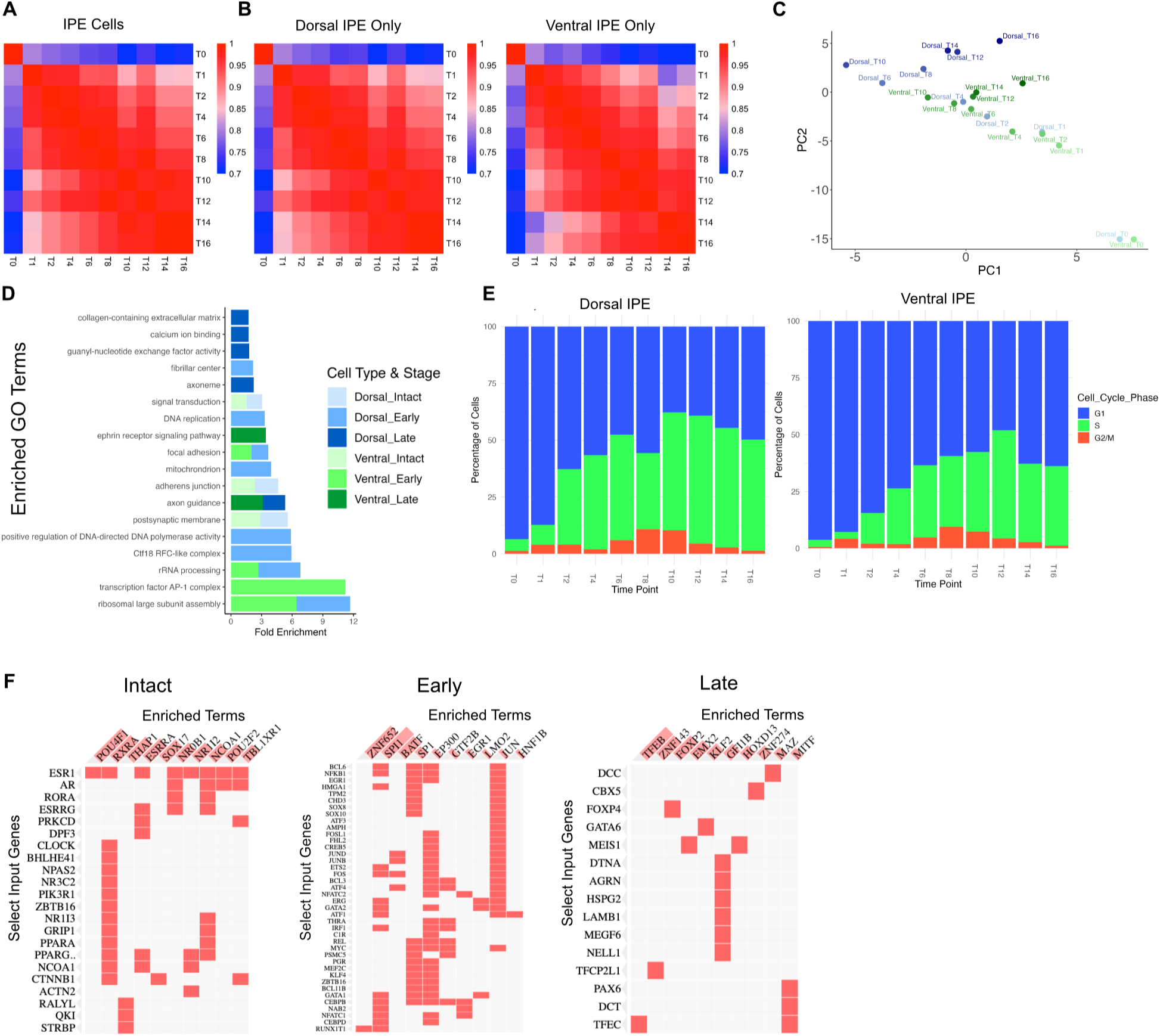
Overview of gene expression changes in IPE cells during regeneration. A) Pearson correlation heatmap for IPE cells across timepoints. B) Pearson correlation heatmaps for Dorsal and Ventral IPE cells across timepoints. C) PCA analysis with Dorsal (blue gradient) and Ventral (green gradient) IPE cells at each timepoint. D) Selected enriched GO Terms from Dorsal (blue) and Ventral (green) IPE cells by stage. E) Cell cycle analysis of both Dorsal and Ventral IPE. G1 in blue, S in green, G2/M in red. F) Predicted transcriptional factors for Dorsal IPE cells by stage.

Overall, our experimental design allowed the direct correlation between molecular and histological stages and provided important clues on how gene expression changed in IPE cells during lens regeneration. IPE cells had a strong neuronal identity and utilized the neuronal machinery for cellular functions like cell movement. Regeneration-competent dorsal IPE cells exhibited increased cell cycle and gene expression dynamics compared to ventral IPE cells. At the genetic level, it was predicted that gene expression regulation was shifted from hormone receptors in intact, to chromatin remodelers early, to developmental genes late in regeneration. These changes reflected the stem cell-like state of IPE cells assumed early during dedifferentiation^26–28^, and fate switch and lineage commitment at the end of the IPE-to-LEC reprogramming process.

### Intact newt iris contains multiple distinct IPE subpopulations

One of the outstanding questions in the field is whether the newt iris contains multiple IPE subpopulations. To address this, neuroepithelium-derived PCE, NPCE, and IPE cells from the intact sample were selected and subclustered (Figure 3A). This analysis revealed 16 putative subpopulations: Five PCE, three NPCE, six IPE, one cluster of ciliary muscle cells, and one composed of cycling cells (Figure 3B). NPR1 expression was found in all IPE subpopulations while LTBP2, the previously identified marker for IPE cells from all timepoints, appeared differentially expressed (Figure 3C and Figure 1E). IPE cells during lens regeneration change their fate, so it was expected that IPE cells may not be defined using a single marker for all molecular stages and timepoints. While LTBP2 remains a good overall marker for IPE cells (Figure 1), some clusters in the intact iris show relative low expression (Figure 3C; LTBP2) and it is also almost absent in converted LECs (discussed later). Cells with low LTBP2 expression (LTBP2^low^) had increased levels of CHRM3 (CHRM3^high^) indicating that these two genes were good markers for identifying certain subpopulations in the intact iris (Figure 3C; LTBP2 and CHRM3).

**Figure 3:**
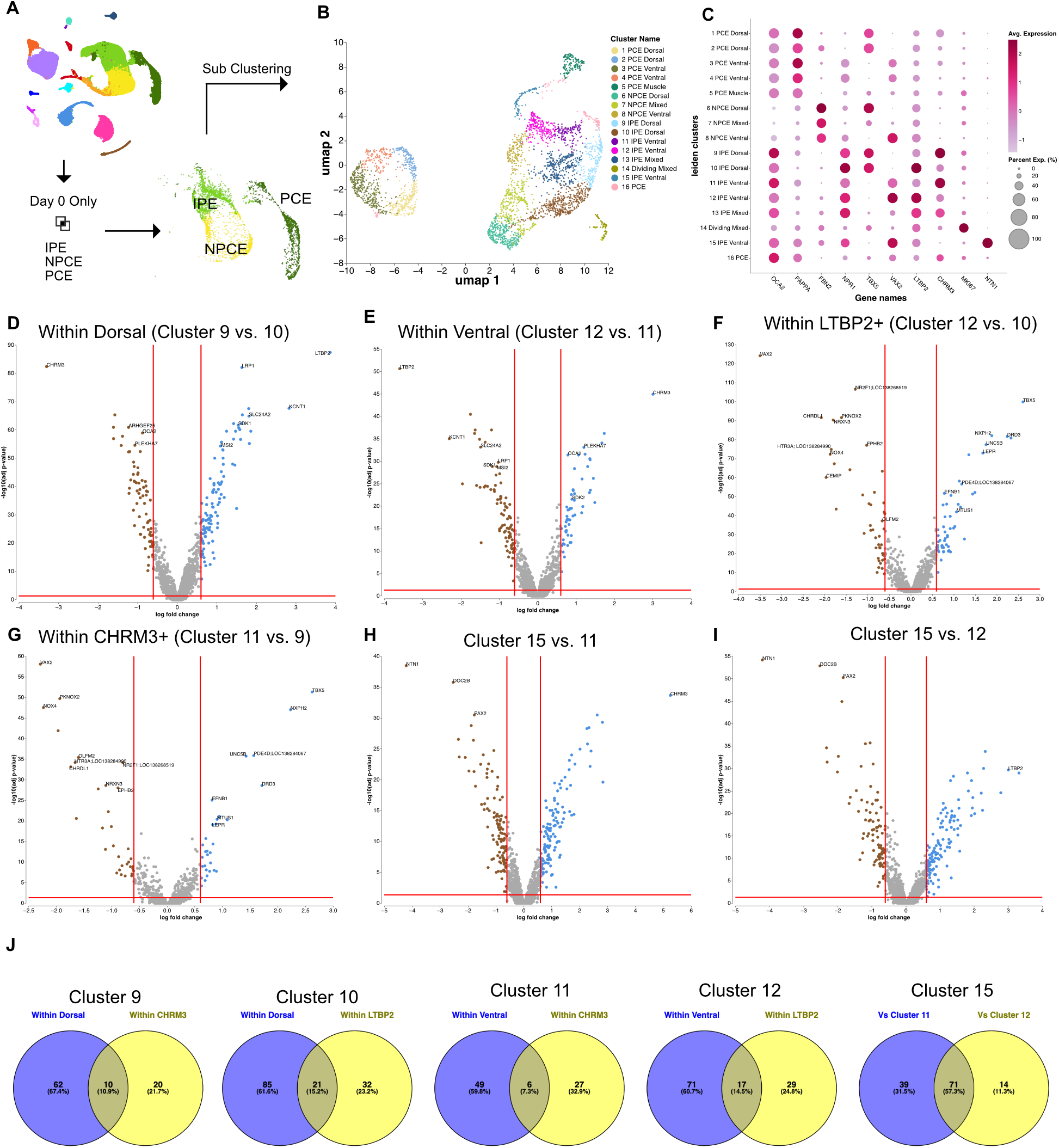
Identification of subpopulations in the intact iris epithelium. A) Experimental design. B) UMAP of intact IPE, NPCE, and PCE cells colored by clusters. C) Dot plot of marker genes for each cluster. D-I) Cluster comparisons using volcano plots with cutoffs adj p-value <0.05, logFC >0.6, and logFC <-0.6. D) Dorsal IPE clusters 9 and 10. E) Ventral IPE clusters 12 and 11. F) LTBP2+ clusters 12 and 10. G) CHRM3+ clusters 11 and 9. H) Ventral IPE cluster 11 and NTN1+ cluster 15. I) Ventral IPE cluster 12 and NTN1+ cluster 15. J) Unique and shared genes across clusters.

To characterize the subpopulations in detail, the following marker genes were used: TBX5 (dorsal marker), VAX2 (ventral marker), PAPPA (PCE marker), FBN2 (NPCE marker), OCA2 (pigment marker), NPR1 (IPE marker), LTBP2 (IPE subpopulation marker), and CHRM3 (IPE subpopulation marker) (Figure S3). There were four PAPPA+ FBN2-LTBP2-PCE subpopulations, two of which were TBX5+ VAX2- in the dorsal iris (Figure 3B and 3C; Clusters 1 and 2), and two VAX2+ TBX5- in the ventral iris (Figure 3B and 3C; Clusters 3 and 4). Cluster 16 contained OCA2+ pigmented cells with mixed dorsoventral identity and an expression profile resembling that of PCE but with relatively lower PAPPA levels. There were three FBN2+ PAPPA-LTBP2^low^ NPCE subpopulations, one of which was TBX5+ VAX2- in the dorsal iris (Figure 3B and 3C, Cluster 6), one VAX2+ TBX5- in the ventral iris (Figure 3B and 3C; Cluster 8), and one with mixed dorsoventral identity (Figure 3B and 3C; Cluster 7). A small number of MKI67+ cycling cells from IPE, NPCE, and PCE were found in cluster 14 (Figure 3B and 3C). Cluster 5 strongly expressed genes found in ciliary muscle like MYH11 and TPM2^17^ (Table S6). There were six putative NPR1+ OCA2+ FBN2-PAPPA-IPE subpopulations, two of which were TBX5+ VAX2- in the dorsal iris (Figure 3B and 3C, Clusters 9 and 10), three VAX2+ TBX5- in the ventral iris (Clusters 11, 12, 15), and one of mixed dorsoventral identity (Cluster 13). IPE clusters 9, 10, 11, 12, and 15 were systematically compared to identify unique marker genes and functions (Figure 3D-I). CHRM3 and LTBP2 were identified as the most differentially expressed when the two TBX5+ dorsal IPE clusters 9 and 10 (Figure 3D) and the two VAX2+ ventral IPE clusters 11 and 12 (Figure 3E) were separately compared. This indicated a two binary cellular identity where IPE subpopulations can be characterized using the dorsoventral markers TBX5 or VAX2, and CHRM3 or LTBP2. CHRM3 and LTBP2 may define a different positional axis for IPE cells like the anteroposterior or proximodistal axis in the iris bilayer. However, a clear association between the enriched genes found in these clusters and preexisting single cell datasets with position axis could not be identified^17^. CHRM3^high^ IPE cells had higher levels of the pigment marker OCA2, the structural component of zonula adherens junctions PLEKHA7, the growth cone inhibitor SEMA3D, and the stress fiber organizer ARHGEF25 (Figure 3D; brown and Figure 3E; blue). LTBP2^high^ IPE cells had higher levels of the neuronal-associated genes KCNT1, LRP1, SLC24A2, MSI2, and SDK1 (Figure 3D; blue and Figure 3E; brown). To further characterize the LTBP2^high^ and CHRM3^high^ subpopulations, we compared the LTBP2^high^ IPE clusters 10 and 12 (Figure 3F) and the CHRM3^high^ IPE clusters 9 and 11 (Figure 3G). As expected, the dorsoventral markers TBX5 and VAX2 were the most differentially expressed genes for both comparisons. In addition, TBX5+ IPE cells were also enriched for NXPH2, UNC5B, PDE4D, DRD3, EFNB1, MTUS1, and LEPR. VAX2+ IPE cells were enriched for PKNOX2, NOX4, OLFM2, HTR3A, NR2F1, CHRDL1, EPHB2, and NRXN3 (Figure 3F and 3G). Cluster 15 which represented the third VAX2+ ventral IPE subpopulation overexpressed NTN1, PAX2, and DOC2B, compared to the other ventral subpopulations (Figure 3H and 3I). VAX2+ NTN1+ cluster 15 appeared to represent a subpopulation only residing in the ventral iris with likely functions in the suturing of the iris^29^. To identify potentially overrepresented and unique genes for each of these subpopulations, gene expression comparisons between the enriched genes from each cluster were performed (Figure 3J and Table 1). The analysis revealed the cell surface proteins cadherin CDH12 and neural cell adhesion protein NCAM2 enriched in the CHRM3^high^ dorsal IPE cluster 9, and cadherin CDH4 and gap junction protein GJD2 enriched in the LTBP2^high^ dorsal IPE cluster 10. These antigens could be used in the future to separate the two subpopulations and conduct studies related to their regenerative ability.

**Table 1:**
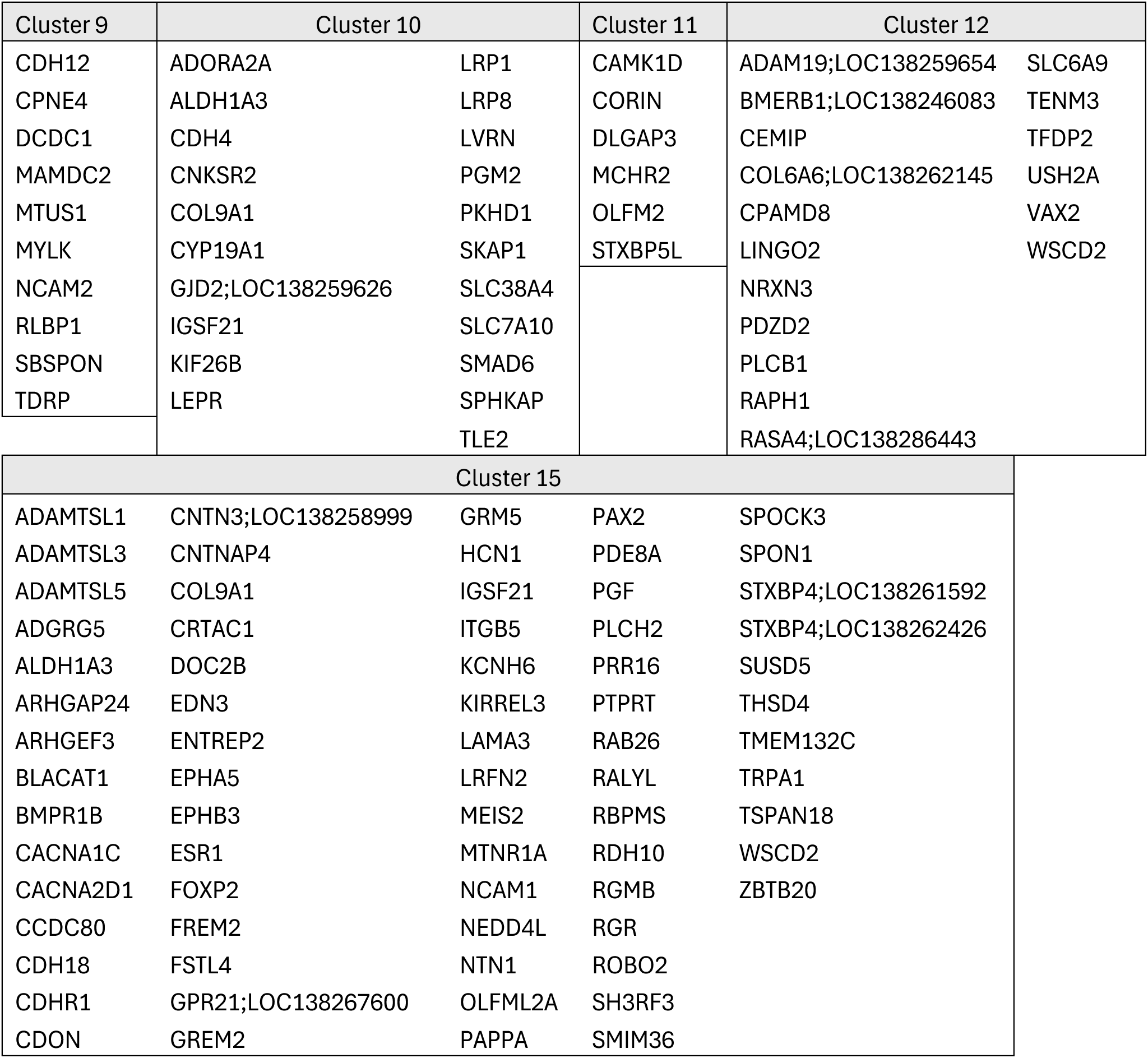
Enriched genes in selected Dorsal IPE clusters.

Overall, the analysis of the intact iris has identified multiple different subpopulations and gene markers that define them. In addition to the previously known dorsoventral markers TBX5 and VAX2, CHRM3, LTBP2, and NTN1 were also identified as markers for different IPE subpopulations. Candidate genes including cell surface antigens were also identified and could be used for the functional characterization of each subpopulation.

### Dorsal IPE cellular states, fates, and functions during newt lens regeneration

Following the identification and characterization of IPE subpopulations in the intact iris, we investigated the cellular states that they assumed during their conversion to LECs. To accomplish this, dorsal IPE cells were subclustered from all timepoints and revealed 16 clusters repressing putative states acquired during lens regeneration (Figure 4A). The clusters did not correlate with specific timepoints except for intact IPE cells primarily found in cluster 4 (Figure 4A; cyan and 4B; black). Clusters showed distinct transcriptomic profiles and revealed the cellular heterogeneity during the IPE-to-LEC conversion (Figure 4C). A timecourse-dependent list of genes is also provided in Figure S4A. Expression analysis of cluster-enriched genes indicated that the late-stage cluster 2 contained presumptive converted LECs expressing the transcriptional factor prospero homeobox protein 1, PROX1 (Figure 4D; PROX1). In the eye, PROX1 is specifically expressed by LECs and lens fibers^30,31^. LEC cluster 2 was also enriched with lens-associated genes like the gap junction GJA8 a structural component of the lens^32^ (Table S7; cluster 2). Cluster 4, which contained intact IPE, highly expressed CHRM3, NPR1, and LTBP2 (Figure 4D). CHRM3 and NPR1 appeared mostly in cluster 4 while LTBP2 was expressed in most clusters except in LEC cluster 2 (Figure 4D). These data indicated that certain IPE markers are quickly lost during regeneration and further strengthened the notion that IPE cells can be identified by LTBP2 expression throughout reprogramming up until their differentiation to LECs. Frequency analysis across timepoints revealed that certain clusters were predominantly found in certain stages like cluster 4 in intact, cluster 1 early and cluster 2 late during regeneration (Figure 4E and 4F). GLM slope analysis indicated that during regeneration intact cluster 4 reduced in frequency while regeneration clusters 2, 11, 12, and 14 increased their presence (Figure S4B). To identify whether the IPE clusters corresponded to different cellular states with potential specific functions, Gene Ontology enrichment analysis was used. Intact cluster 4 was found enriched in melanin biosynthesis genes, a function correlating with them being pigmented and the fact that IPE cells are known to shed their pigment granules during lens regeneration^11^ (Figure 4G; cyan). Early-stage cluster 1 was enriched in genes associated with DNA replication indicating that these cells were in the S phase (Figure 4G; orange). Cluster 9, which appeared in both early and late stages, was enriched in genes that regulated chromosome segregation which indicated that these cells were in the M phase of the cell cycle (Figure 4G; dark blue). The temporal appearance of clusters 1 and 9 correlated with the normal sequence of the cell cycle and with our cell cycle analysis for the respective timepoints (Figure 2E). Late-stage LEC cluster 2 was enriched in extracellular matrix organization and cell adhesion, two functions closely linked to developing a lens vesicle with a capsule (Figure 4G; yellow). Clusters 8 appeared in both early and late stages of regeneration and was enriched for functions related to protein folding chaperone, intermediate filament, and extracellular exosome (Figure 4G; purple). Cluster 8 overexpressed NCL which codes for Nucleolin a protein involved in cell growth (Figure 4C and Table S7). IPE cells grow in preparation for the cell cycle, thus, cluster 8 likely represented cells at the G1 or G2 phases (Figure 1B). Cluster 13 also appeared in both early and late stages of regeneration and was enriched for keratin filament and collagen-containing extracellular matrix (Figure 4G; blue). These cells overexpressed KRT8 and CLDN5 which indicated IPE cells likely retaining their epithelial fate and structure and did not actively undergo reprogramming (Figure 4C and Table S7). Cluster 10 appeared at the very early and late timepoints of regeneration. This cluster was enriched for positive regulation of defense response to virus by host and overexpressed genes like the anti-viral HERC5. This indicated that these cells were under stress likely due to the inflammatory response triggered following lens removal. Clusters 0, 3, 5, 6, 7, 11, 12, 14, and 15 did not have any enriched functions. Gene expression enrichment analysis indicated that early-stage cluster 6 likely included migratory cells overexpressing the focal adhesion cadherin CDH11. Late-stage clusters 11, 12, 14, and 15 included IPE cells that overexpressed neural, pigment, and guidance genes like the neurotransmitter regulator DBH and ion channel CACNA1C in 11, phototransduction gene RGS9 and netrin receptor DCC in 12, melanosome regulator HPS5 in 14, and the adherens junction cadherin CDH12 in 15. These gene signatures indicated that IPE cells in these clusters did not actively participate in the reprogramming process and retained an IPE fate. To further analyze the association between gene expression and a particular cell state or function, gene expression correlation analysis was performed for all genes expressed by the clusters. Figure 4G lists the top 4 positively or negatively correlated gene pairs in each cluster. The data revealed six consistent expression signatures: (a) LTBP2 was positively regulated with MECOM and LRP1 in clusters 1, 4, 8, and 9, and inversely correlated with CIRBP, PMEPA1, GATAD2B, FBXO27, EIF2S2, CHRM3, MKI67, and TACR3, in clusters 0, 1, 4, 8, 9, and 15. (b) CHRM3 was positively correlated with SLC6A13, SGCD and SMTNL2 in clusters 7, 14, 15, and inversely correlated with MMP13, SYT7, RUNX1, LTBP2, ESYT1, IQGAP2, HSPG2, NR6A1, and COL26A1 in clusters 3, 4, 5, 6, 7, 10, 13, 14, and 15. (c) Growth regulators HMGB2, NCL, TRIM69, SET, and HNRNPAB were positively correlated in clusters 1, 8, and 9, and inversely correlated with MECOM, BCAS3, and MACROD2 in cluster 11. (d) Cell cycle regulators CCNB3, FAM167A, ASPM, AURKA, KIF11, E2F7, PBK, DTL, NDC80, CDK1, ESCO2, KCNK2, ZWINT, CCNB1, RSPO3, UHRF1, E2F2, SUV39H1, DEPDC1, PRR11, ECT2, ORC6, HASPIN, SAMD8, CDC6, RAD17, BORA, CEP55, and PCNA were positively correlated in clusters 3, 4, 5, 7, 10, 11, 12, 14. (e) Cell surface and extracellular matrix genes GLDN, SORCS1, NRCAM, IGFBP5, COL4A5, COL4A6 were positively correlated in clusters 2, and 6, and inversely correlated with FN1 in cluster 2. (f) NR6A1, IQGAP2, MMP13, HSPG2, ADAMTS12, and COL26A1 were positively correlated in clusters 3, 10, 13, and 15, and inversely correlated with ABCB5, LVRN, PDZRN3, SLC6A13, CHRM3, ADGRG1, and CYP2A13 in clusters 1, 3, 5, 6, 7, 10, 12, 13, 14, and 15. Collectively, these data underscored the importance of gene expression regulation and coordination in cell cycle entry, cell fate determination, extracellular matrix remodeling and signal transduction during lens regeneration.

**Figure 4:**
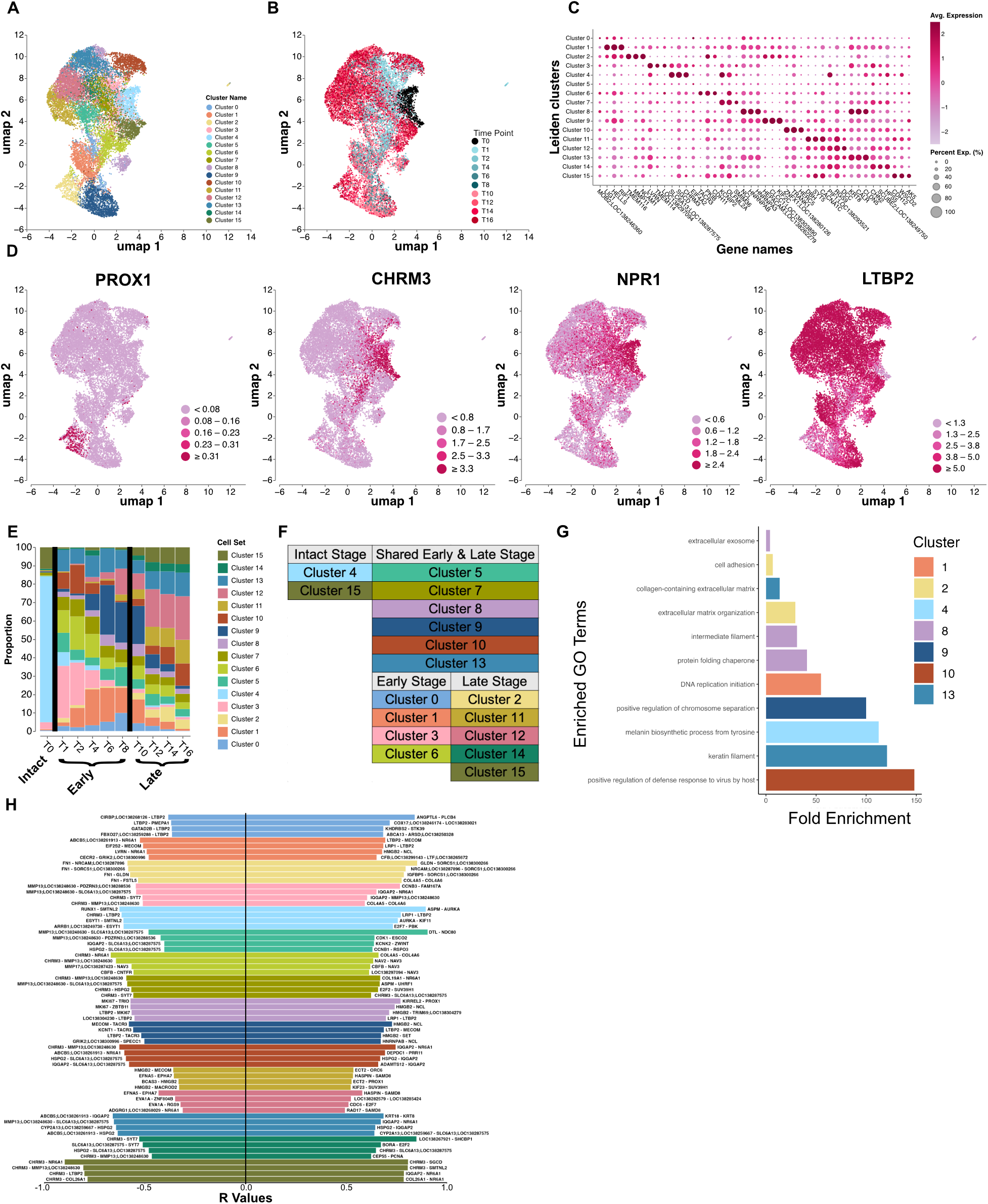
Identification of dorsal IPE states during lens regeneration. A) UMAP of Dorsal IPE cells colored by cluster. B) UMAP of Dorsal IPE cells colored by timepoint. C) Dot plot of top three marker genes per cluster. D) UMAPs of Dorsal IPE cells colored by PROX1, CHRM3, NPR1, and LTBP2 expression. E) Frequency plot of clusters across timepoints. Stages indicated at the bottom. F) Summary of cluster distribution across stages. G) Selected enriched GO Terms from clusters. H) Top 4 positively and negatively correlated gene pairs per cluster.

Overall, following lens removal, IPE cells down-regulate intact marker genes like CHRM3 and transition from the intact adherens junction-based structure to a more mobile focal adhesion-dependent state. IPE cells enter the cell cycle and can be identified at every phase with distinct expression profiles. Late-stage IPE clusters were found bifurcated with the presumably converted LECs expressing the lens transcriptional factor PROX1 among other lens-associated genes while the rest strongly expressing neuronal-associated genes. These profiles reflected cells committing to a lens fate or retaining the IPE fate, a function that ensures the integrity of the iris following regeneration. Finally, gene expression correlation analysis revealed genes associated with certain IPE functions like cell cycle and extracellular matrix regulation, as well as the potential antagonistic roles of CHRM3 and LTBP2.

### Cellular trajectories reveal the molecular blueprint of IPE-to-LEC reprogramming

Our experimental design encompassed multiple timepoints in short sequence during lens regeneration to capture the cellular heterogeneity of IPE cells as they reprogram into LECs. While our analysis in Figure 4 captured clusters of IPE cells representing different cellular states and functions assumed during regeneration, it lacked a direct analysis of how cellular trajectories and fates changed over time. To accomplish that, Monocle3 was used in dorsal IPE cells to generate embedded predicted cellular trajectories. PROX1+ LECs appeared at the edge of the trajectories (Figure S5A). Two main cellular paths were revealed, one of which highly correlated with the actual timepoint sampling while the second was represented by late-stage timepoints (Figure S5B). The almost identical alignment of the unsupervised pseudotime embedding with the actual timecourse was intriguing and potentially reflected the reprogramming process, thus, cells from this path were selected for further downstream analysis (Figure 5A and Figure S5C). PROX1+ LECs were represented by Days 14 and 16 and were selected as the end of the trajectory (Figure 5B). Pseudotime was computed in 42 different nodes (Figure 5C) and used to rank and order gene expression across the trajectory (Figure 5D). In line with previous observations, CHRM3 was highly expressed in nodes early in the trajectory and was down-regulated thereafter. At later timepoints, there was up-regulation of cell cycle regulators like SMC2 and KIF14, DNA damage response proteins like BRCA1 and FANCI, and cell signaling transduction regulators like MAP2K6. In addition to how genes changed overtime, expression regulation was also investigated to reveal potential regulatory networks involved in the IPE-to-LEC cell fate switch. To accomplish that, genes were first divided into different modules based on their expression pattern across the trajectory (Figure S5D). For instance, gene module 3 contained CHRM3 and had a pattern with high expression early in the trajectory and decreased with time. Gene module 11 contained MAP2K6 which had low expression early but increased with time. Gene module 12 contained genes like SYT7 that had high expression during reprogramming. Lastly, Gene module 10 contained PROX1 with high expression only at the end of the trajectory (Figure 5E and 5F). Next, Enrichr was used to identify putative reprogramming and transcriptional factors that regulated the expression of genes contained in each module^25^. Early expressed module 3, was regulated by MITF, a known transcriptional factor found in pigmented cells highlighting the IPE fate at the onset of regeneration^33^. Injury-induced module 11 was governed by differentiation, stem cell maintenance and chromatin remodeling factors including GFI1B, SOX2, HDAC2, RNF2, ESRRB, MTA3, and RCOR1. This illustrated the dedifferentiation and reprogramming events necessary for converting IPE cells to LECs. Module 12 was regulated by FOXP2, a transcriptional factor that specifies and organizes epithelial tissues which indicated changes in the structure and fate of IPE cells to accommodate the making of a lens vesicle^34,35^. Lastly, module 10 which contained the majority of PROX1+ LECs, was regulated by cell determination factors including JARID2, LMO4, NR4A2, and TFAP2A. Interestingly, NR4A2 is a differentiation factor for mesodiencephalic cells potentially indicating the depth of dedifferentiation of IPE cells during reprogramming^36^. In addition, TFAP2A is a transcriptional factor known to regulate early morphogenesis of the lens vesicle during development^37,38^. Taken together, IPE-to-LEC conversion was regulated by MITF in IPE cells, SOX2 and HDAC2 during dedifferentiation, JARID2 and NR4A2 during fate commitment, and TFAP2A and PROX1 in converted LECs.

**Figure 5:**
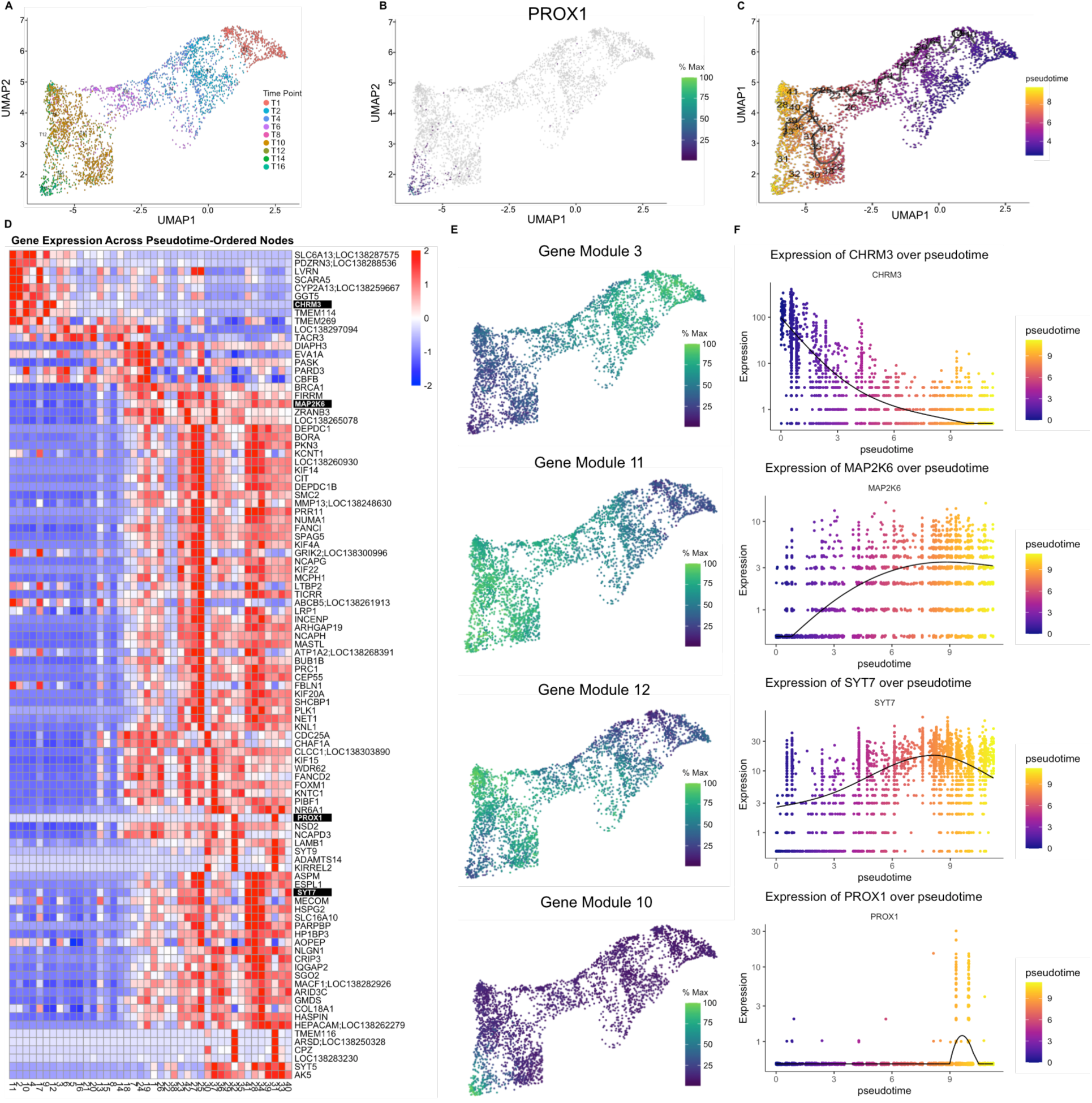
Cellular trajectory analysis of dorsal IPE cells during lens regeneration. A) Monocle3-derived UMAP of Dorsal IPE cells colored by timepoint. B) UMAP colored for PROX1 expression. C) UMAP with branches and node labels related to pseudotime. Pseudotime indicated in purple for early and yellow for late points. D) Heatmap with the top three marker genes of each node ordered by pseudotime. E) Selected modules 3, 11, 12, and 10 for different gene expression patterns across pseudotime. F) Linear expression plots of selected genes CHRM3, MAP2K6, SYT7, and PROX1 representing the selected modules from E.

Overall, cellular trajectory combined with gene expression analysis has revealed new insights into IPE-to-LEC reprogramming. Our datasets track gene expression as IPE cells convert to LECs and predict how major transitions are regulated at the genetic level.

### Tissue-resident macrophages polarize from M1 to M2 during lens regeneration

The vertebrate iris contains many different cell types most of which were captured by our scRNAseq experiment. In addition to IPE cells, cells from the adjacent NPCE and PCE, cells residing in the stroma including melanocytes, iridophores, endothelial cells, and fibroblasts, cells from the blood including B cells, T cells and macrophages were also captured and analyzed (Figure 1). To identify potential roles of these cells during lens regeneration, ligand-receptor interaction analysis was used (Figure 6A). NPCE, IPE, fibroblasts, PCE, and macrophages accounted for the majority of the signaling. The most significant interactions were presented in Figure 6B and Figure S6A. These data indicated that dorsal IPE cells received CLCF1 from fibroblasts, MDK, TGFB1, GDF11 from NPCE, MDK and GRN from PCE, and GRN and TGFB1 from macrophages. Past studies have shown that macrophages interact with IPE cells, engulf pigment granules, increase their numbers, and potentially provide signaling molecules during lens regeneration^10–12^. To characterize macrophages in greater detail, the CSF1R+ macrophage cluster from Figure 1 was subclustered and revealed 16 subpopulations representing different putative states assumed during regeneration (Figure 6C). Clusters appeared to correlate with regeneration stages (Figure 6D and Figure S6B). Gene expression analysis revealed distinct timepoint-dependent profiles divided into three stages, intact, early (Days 1-8), and late (Days 10-16), resembling the regeneration stages of IPE (Figure 6E). To identify different macrophage subtypes, gene expression analysis of known markers was performed. Macrophages were found positive for CD68, a tissue resident marker. This indicated that during regeneration macrophages are homed to the IPE from within the eye (Figure 6F). The ligand-receptor data suggested that this function was likely due to CSF1 secretion from IPE cells, a ligand known to interact with the CSF1R receptor in macrophages and activate them^39^ (Figure 6B). Gene expression analysis also revealed that during the early-stage macrophages highly expressed IL18, an inflammatory molecule associated with the M1 macrophage subtype^40^ (Figure 6G). During the late-stage, macrophages overexpressed ADGRE1 a molecule associated with the M2 macrophage subtype^41^ (Figure 6H). Late-stage macrophages also expressed the proliferation marker MKI67 (Figure 6I) and were found to enter the S and G2/M phases of the cell cycle (Figure 6J), an event also associated with the M2 subtype^42,43^.

**Figure 6:**
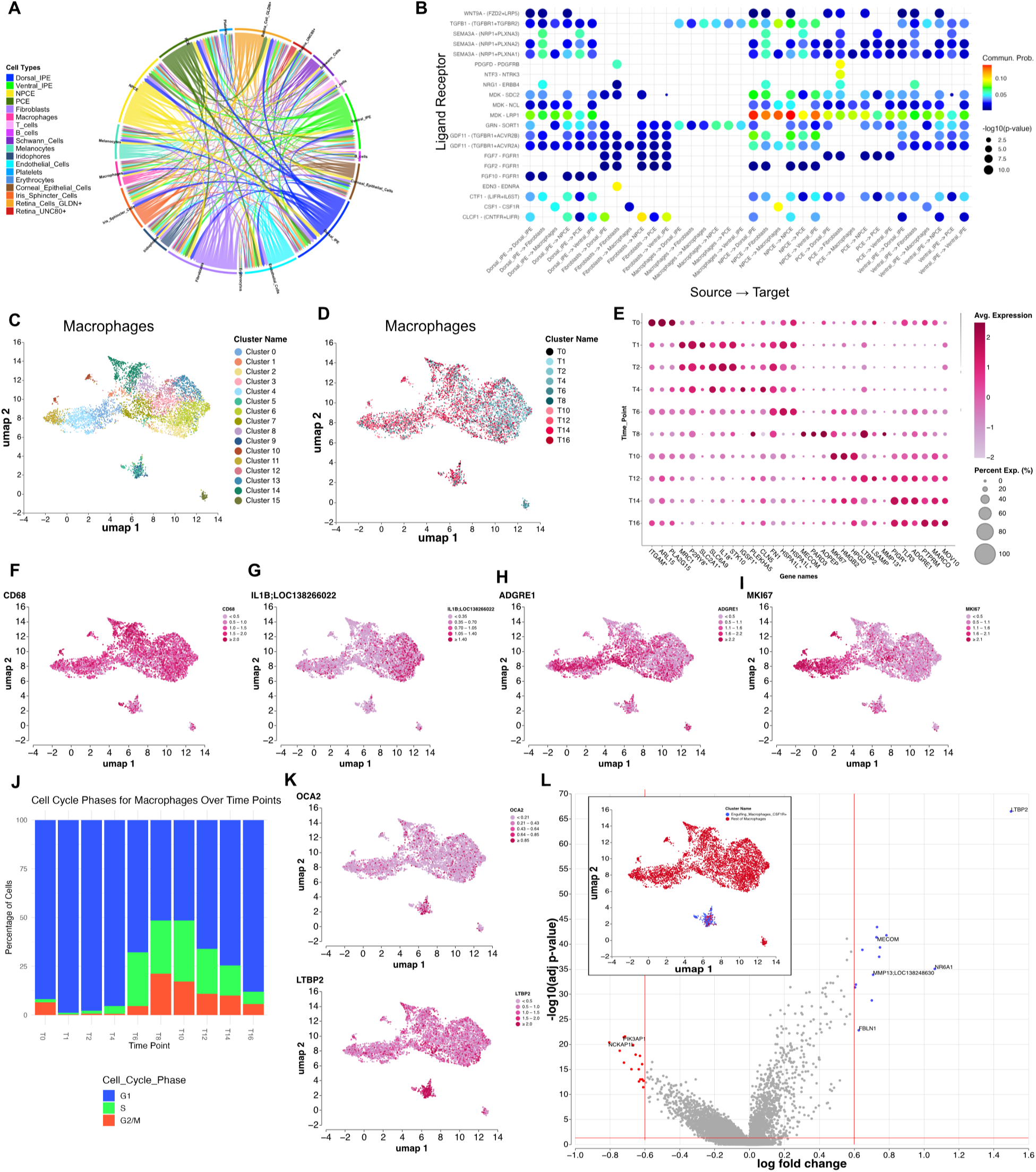
Cell-cell Interactions and macrophage heterogeneity during lens regeneration. A) Chord diagram depicting all ligand-receptor interactions per cell type. B) Bubbleplot depicting selected significant interactions. C-I) Data from the macrophage only subset C) UMAP of macrophages colored by cluster. D) UMAP of macrophages colored by timepoint. E) Dot plot with the top three marker genes per cluster. F) UMAP of CD68 expression. G) UMAP of ADGRE1 expression. H) UMAP of IL1B;LOC138266022 expression. I) UMAP of MKI67 expression. J) Cell cycle analysis of macrophages over time. G1 in blue, S in green, G2/M in red. K) Volcano plot of the “Engulfing macrophage CSF1R+” subset versus all other macrophages. UMAP of the groups compared is embedded.

**Figure 7:**
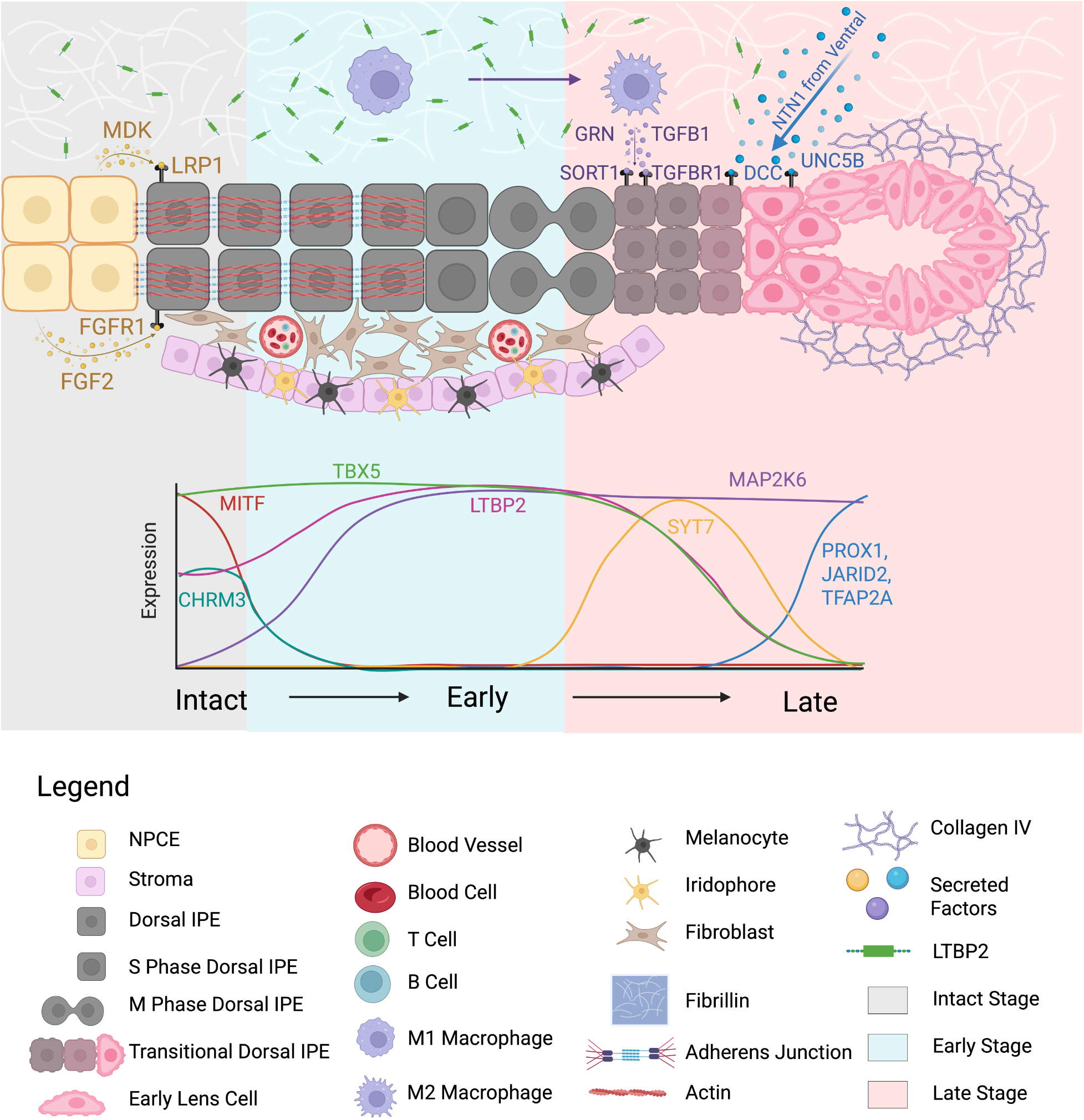
Graphical Summary. Cartoon summarizing major findings. Changes of selected cell types, signaling pathways, and genes during lens regeneration are displayed.

Our subclustering analysis also revealed two subpopulations of macrophages that were enriched with IPE-related genes including the pigment marker OCA2 and the IPE marker LTBP2 (Figure 6C and 6K). Gene expression comparison of these two clusters and the rest of macrophages revealed additional IPE-related genes including MECOM, NR6A1, MMP13, and FBLN1 (Figure 6L). To exclude potential doublets that were not filtered out, a secondary doublet analysis was performed and revealed canonical number of reads per cell with no signs of incomplete cell dissociation between IPE cells and macrophages (Figure S6C), indicating that these genes were derived from the macrophages. IPE cells are known to secrete melanin granules which are engulfed by adjacent macrophages^11^. Our data indicated that IPE cells may actively secrete other cell contents including mRNA which were also engulfed by macrophages. Our scRNAseq experiment likely captured engulfed IPE-derived mRNA in macrophages before their eventual degradation. Secretion of cell contents including membrane-bound proteins and mRNA by IPE cells could accelerate their reprogramming since IPE-related transcripts and proteins are not needed and will be replaced by that of LECs.

Overall, IPE cells are a signaling hub to and from adjacent cells including the secretion of CSF1 for the activation of macrophages. In turn, macrophages secret TGFB1 and GRN and engulf IPE contents potentially including mRNA. Macrophages were identified as CD68+ tissue resident and polarized from inflammatory IL18+ M1 to anti-inflammatory ADGRE1+ M2 type during regeneration. The M1-to-M2 polarization of macrophages was highly correlated with the reprogramming stages of IPE indicating dependency. The overall increase in macrophage number adjacent to the IPE can be attributed to proliferation, especially at later timepoints, and attraction by guiding molecules like the IPE-secreted CSF1.

## Discussion

Here we present the first single cell Atlas of the newt iris during Wolffian lens regeneration. It includes 80,425 deeply sequenced cells from intact and nine timepoints following lens removal iris tissue. We provide an extensive yet compelling analysis of the data starting with (a) identifying and characterizing different cell types in the iris and adjacent tissues, (b) revealing global changes in gene expression for all cell types, (c) uncovering new previously uncharacterized IPE subpopulations in the intact iris, (d) identifying and characterizing different functional IPE states assumed during lens regeneration, (e) interrogating gene expression and genetic regulation during IPE-to-LEC reprogramming with pseudotime trajectory analysis, and (f) revealing new roles and functions for non-IPE cells including new roles and functions for macrophages. Overall, this atlas is a comprehensive resource for the tissue regeneration scientific community and can be used to generate hypothesis and base future experimentation to further understand how newts regenerate their eye tissues.

During newt IPE-to-LEC reprogramming, IPE cells change their cellular appearance from completely pigmented to completely transparent. For each of our timepoints used in making the scRNAseq atlas, sibling newts were also collected to provide matched histological data on the iris (Figure 1A and 1B). This provided the ability to directly compare and correlate gene expression and histological changes from similar newts. Our experimental design also intentionally enriched for the regeneration-competent dorsal iris since the main goal was to uncover the mechanism of IPE-to-LEC reprogramming.

However, lens regeneration-incompetent ventral iris was also included to potentially identify new cells and functions and contribute to the overall completeness of this atlas. The dorsal to ventral cell ratio in the atlas reflected the performed extractions (Figure S2B). The iris is composed of the stroma in the anterior side which contains a variety of cells most of which were captured by our scRNAseq experiment including melanocytes, iridophores, fibroblasts, iris sphincter cells, Schwann cells, endothelium, and blood-derived B cells, T cells, nucleated erythrocytes, and platelets. Blood-derived cells were retained in the iris vasculature during extraction with no evidence that they play a role during lens regeneration (Figure S2A). The captured macrophages were tissue resident, mainly derived from the eye itself, likely in direct or close contact with the IPE cells^11^ (Figure 6 and Figure S6). Technically, the extraction of pure iris samples is challenging. The iris ring near the pupil transitions to the NPCE and PCE ciliary tissues which were also represented in the atlas (Figure 1D). Following lens removal, the eye collapses and retina tissue was observed near the iris during extractions. Lastly, during the peeling of the iris from the cornea, especially in early timepoints when the cornea was undergoing a healing process itself, cornea cells were likely attached to the iris and included during single cell isolation. Each of our samples contained numerous stringently extracted irises from different sibling newts to generate an atlas mainly represented by IPE cells despite these challenges.

We identified three gene expression-based stages: intact, early (Days 1-8), and late (Days 10-16). Following lens removal, cell response measured as a relative change in gene expression, followed a proximodistal gradient along the iris with the most observed in IPE cells followed by NPCE, PCE, and retina (Figure S2A). Ventral IPE cells also responded to injury in a similar manner. However, there was a significant delay in progression with late-stage ventral resembling that of 4dpl early-stage dorsal IPE cells (Figure 2C). Past studies have attributed the ability of dorsal IPE cells to successfully reprogram to a LEC fate to their speed in entering the cell cycle and reprogramming^20,21^. This is largely reflected in our gene expression analysis with dorsal IPE cells responding the most following lens removal (Figure 2A). Two hypotheses for this response includes the expression of the protein tissue factor in the dorsal iris and the secretion of FGF ligands by macrophages around the dorsal iris^12,21^. However, our scRNAseq experiment does not directly support them with the tissue factor gene F3 showing no expression in IPE cells (Table S2) and FGF ligands only provided by the NPCE and fibroblasts while macrophages found secreting TGFB1 and GRN (Figure 6B). Despite the deep sequencing of the cells in this atlas it is possible that these genes were below the detectable levels for the respective cell types. Our data corroborated histological evidence showing an increase in the number of macrophages associated with the iris during regeneration^11^ (Figure 6J). Our detailed single cell analysis of macrophages revealed for the first time that they are predominantly CD68+ tissue resident, suggesting that the eye maintains its immune privileged state during lens regeneration^44^. In addition, we identified for the first time that macrophage polarize from the inflammatory M1 early to the anti-inflammatory M2 subtype late during regeneration (Figure 6E). The M1-to-M2 macrophage polarization was highly correlated with the IPE-to-LEC reprogramming which may indicate dependency for successful regeneration. It is unclear whether macrophages, IPE cells, or a different cell type was responsible in regulating these transitions. The correlation between the processes could also have emerged by the temporal presence of a factor in the vitreous humor that affected cellular responses at similar timepoints. Overall, these data underscored the importance of the regenerative microenvironment in regulating and promoting regeneration.

Past studies have shown that newt dorsal IPE cells regenerate lens tissue in a variety of environments including eyes of lens regeneration incompetent species^3,45,46^. This strongly indicates that this ability is innate in newt dorsal IPE cells. Identifying and characterizing the very IPE cells capable of LEC reprogramming would further their study including their direct comparison with IPE cells from other organisms. The use of bulk RNA sequencing had previously identified TBX5 as a marker gene for the regeneration competent dorsal iris and VAX2 as a marker gene for the incompetent ventral iris^7^. Our scRNAseq data further increased our understanding of the IPE cell heterogeneity in the intact iris with three additional marker genes, CHRM3, LTBP2, and NTN1 (Figure 3G). NTN1+ IPE cells were found only in the ventral iris, and this subpopulation can be defined as VAX2+ TBX5-NTN1+. Interestingly, this subpopulation also expressed the transcriptional factor PAX2, a gene known to cause coloboma, a condition usually associated with the incomplete suturing of the pupil at the ventral side^29^. The other marker genes, the dorsoventral-associated TBX5 and VAX2, together with CHRM3 and LTBP2 formed a two binary system that specified four different IPE subpopulations: (1) TBX5+ VAX2-CHRM3^high^ LTBP2^low^, (2) TBX5+ VAX2-CHRM3^low^ LTBP2^high^, (3) TBX5-VAX2+ CHRM3^high^ LTBP2 ^low^, and (4) TBX5-VAX2+ CHRM3^low^ LTBP2^high^. Gene expression correlation data also strongly supported the bifurcation of CHRM3 and LTBP2 which appeared as top negatively correlated pairs (Figure 4H). This binary arrangement along the dorsoventral axis for CHRM3 and LTBP2 likely indicates a different positional axis, either the anteroposterior within the epithelial bilayer or the apical/basal axis. CHRM3^high^ cells overexpressed the pigmented gene OCA2 and may be positioned in the more pigmented anterior side of the epithelium (Figure 3G). LTBP2 was highly upregulated during regeneration and LTBP2^high^ cells overexpressed MSI2, a protein found in neural precursor cells, indicating that LTBP2 may identify pro-regeneration IPE cells^47^ (Figure 3G and Figure 4D). LTBP2 could also facilitate a pro-regenerative environment with elastic fibers to support the formation of the lens vesicle at the tip of the dorsal iris^48^. Overall, this atlas has further our understanding on IPE heterogeneity in the intact iris. Future studies will aim to lineage trace, isolate and enrich these subpopulations and test their regenerative capacity.

The collection of multiple timepoints concentrated at the reprogramming stage of lens regeneration provides a molecular blueprint on how dorsal IPE cells convert to LECs. This was reflected to one of the branches of the *in silico*-predicted pseudotime trajectory which highly resembled the sequence of the actual timepoints. We believe that the current and future functional analyses of this trajectory will provide important clues on IPE transdifferentiating. The intact IPE expressed genes associated with adherens junctions, a structure necessary to maintain the integrity of the epithelium bilayer (Figure 2D). MITF was identified to likely regulate this stage (Figure S5E). Following lens removal, intact IPE markers, including CHRM3, SLC6A13, and NPR1, were downregulated relatively fast, within a few days (Figure 4D and Figure 5D). This marked the first instance of IPE dedifferentiation. At the same time, IPE cells started proliferating with all phases of the cell cycle being reflected at their transcriptomic profiles in individual clusters (Figure 2E and 4G). Even when cell cycle-associated genes were regressed from this analysis, we saw little to no change to the clustering of these cells (data not shown). Cell cycle genes were also highly positively correlated which underscores the fundamental link between cell cycle and reprogramming capacity (Figure 4H). Noteworthy, this concept is also described in other cell populations including the ability of retina pigmented epithelial cells and glia to reprogram to retina cells in other model systems^49–51^. IPE cells at this stage also expressed genes associated with focal adhesions potentially indicating epithelial-to-mesenchymal transition and migration (Figure 2D and Figure 4G). This would allow IPE cells to break away from the iris epithelium and begin to form a lens vesicle. We identified that this early stage of regeneration was likely regulated by the FGF signaling effector JUN, Wnt signaling-induced MYC and histone methyltransferase EZH2 (Figure S5E). All three pathways have been previously associated with tissue regeneration and progenitor cell activity in the eye^20,52^. Our atlas identified LTBP2 as an IPE subpopulation marker and a gene that was upregulated during regeneration along with the dorsal marker TBX5. Both LTBP2 and TBX5 were downregulated in PROX1+ LEC cells. These expression profiles indicate a critical cell fate switch between the IPE and LEC lineages. The LEC cluster was also enriched for the early lens specification zinc finger protein CASZ1 and the lens vesicle morphogenesis transcriptional factor TFAP2A (Table S7 and Table S9)^38,53^. Interestingly, TFAP2A was also predicted to regulate the gene module that LECs were part of (Figure S5E). JARID2, a histone methyltransferase activity regulator, was also among the reprogramming factors likely regulating gene expression associated to LEC commitment (Figure S5E). JARID2 is known to impact the timely generation of retina cell types by repressing FOXP1 during development^54^. It is interesting to speculate that the LEC fate is part of such a program with IPE cells dedifferentiating to a neuroepithelial-like progenitor, and the subsequent reprogramming is part of a time-dependent process regulated by FOXP2 and JARID2, two factors appearing sequentially along the IPE-to-LEC trajectory (Figure 2F and Figure S5E). Future studies could potentially affect the activity of genes like JARID2 and allow the generation of retina neurons from the IPE. The LEC cluster also expressed structural components of the lens including the lens gaps junction protein GJA8 and the lens capsule extracellular matrix protein COL4A5 indicating that lens cells were organized soon after LEC commitment and before the lens vesicle is detached from the iris^32,55^ (Figure 1B). Lens crystallin genes were also detectable at low levels (Table S7). IPE cells destined to retain their IPE fate, an important event in preserving the iris structure and the perpetual lens regeneration capacity for these animals, did not downregulate LTBP2 and TBX5 and upregulated guidance genes including the Netrin receptor DCC^2,56,57^ (Figure 4C). Interestingly, dorsal, and not ventral, IPE cells were known to express the Netrin receptor UNC5B, a pattern that was also supported in our Atlas^7^ (Figure 3F and 3G). The netrin ligands appear to originate from the NTN1+ ventral subpopulation and differentially affect dorsal IPE cells depending on the expressed netrin receptors (Figure 3H and 3I). Based on these expression patterns, it appears that netrin signaling regulates IPE migration towards or away from the lens vesicle with DCC and UNC5B potentially marking IPE cells that will be retained in the iris^58,59^. The role of Netrin signaling in IPE cell organization is in addition to the Ephrin signaling in dorsoventral segmentation^24^. Past studies had identified differential expression of Ephrin ligands like EFNB1 in dorsal and receptors like EPHB2 in ventral iris which we also corroborated using scRNAseq (Figure 3F and 3G). Netrin and Ephrin signaling pathways appear to ensure that only certain IPE subpopulations will be present at the tip of the dorsal iris and contribute to lens regeneration.

Overall, this atlas is a rich resource for understanding IPE-to-LEC reprogramming during newt lens regeneration. We provide an extensive yet cohesive analysis addressing the main areas of research with this model and the ability to base hypothesis for future experimentation. We also identified for the first time (1) the cellular heterogeneity of the newt iris and ways to identify each cell type, (2) the presence of multiple IPE subpopulations in the intact iris, (3) the IPE heterogeneity during regeneration including their functional states, (4) a molecular blueprint based on cellular trajectory analysis, (5) cell-cell interactions between IPE and other cells, and (6) the identity of macrophages as tissue-resident and their polarization from M1 to M2 subtype during regeneration.

## Resource availability

Raw and processed data and code will be available upon publication.

## Acknowledgments

We would like to thank members of the Animal Research Office at the University of New Hampshire (UNH) Dr. Dean Elder, Dr. Linnea Morley, and Shelly Morton for excellent animal care. We would also like to thank the University Instrumentation Center for microscopy assistance, the Hubbard Center for Genome Studies for assistance in sequencing and data analysis, and Parse Biosciences for support with their platform. Research reported in this publication was supported by the National Eye Institute of the National Institutes of Health under Award Number R00EY029361 to KS, and by the NH-INBRE program and the Center for Integrated Biomedical and Bioengineering Research (CIBBR) through grants from the National Institute of General Medical Sciences of the National Institutes of Health under Award Numbers P20GM103506 and P20GM113131, respectively. The content is solely the responsibility of the authors and does not necessarily represent the official views of the National Institutes of Health. Research was also supported by UNH’s Summer Teaching Assistant Fellowship to OW.

## Author contributions

Conceptualization, supervision, and funding acquisition done by K.S.; Animal handling done by O.W., K.A., N.F., D.H., K.L., J.L., L.N., J.N., T.R., and B.W.; Surgeries done by K.S.; Isolation and fixation of tissue completed by O.W.; Histological sectioning, staining, and imaging completed by K.A.; Bioinformatics completed by J.S., W.K.T., O.W., and K.S.; Analysis of *in silico* data completed by O.W. with input from K.S.; Visuals generated by O.W.; Text written and edited by K.S. and O.W.

## Materials and Methods

### Animal Procedures

#### Animal Handling

*Pleurodeles waltl* used in this experiment were derived from a single breeding and were raised at room temperature in dechlorinated tap water (Housing water). Animals were 2.5-3.5 months old at the time of extraction. Experiments involving animals were performed in accordance with the UNH Institutional Animal Care & Use Committee Research (IACUC) protocols #220604 and #241003. Animals were euthanized at the end of the experiment by decapitation following anesthesia with buffered 0.1% tricaine.

#### Lens Removal

Animals were anesthetized using buffered 0.1% tricaine. Using a scalpel, a small hole was made at the edge of the cornea. Scissors were used to cut the cornea across its entire length. The lens was retrieved using fine point forceps. Lenses from both eyes were removed. Animals were returned to housing water and resumed normal animal care until tissue extractions. 30 animals had their lenses removed per timepoint, 24 of which were used for iris tissue extraction and 6 for whole eye removals.

#### Iridectomy

Animals were anesthetized, and a hole was made in the sclera just below the plane of the iris and using scissors the anterior segment was removed. Anterior eye segments were removed at 1, 2, 4, 6, 8, 10, 12, 14, 16 days post-lens removal and from intact unoperated newts. Anterior eye segments containing cornea, iris, and adjacent tissues were moved to cold 0.7X PBS without Ca and Mg. Using a scalpel, the iris ring was cleaned from other tissues. For 32 irises (16 newts), the dorsal segment was separated, peeled off the cornea and transferred to a Bovine Serum Albumin-blocked DNA LoBind Eppendorf tube containing cold 0.7X PBS without Ca and Mg. 16 additional irises (8 newts) were collected as a whole containing both dorsal and ventral segments. In total, there were 48 dorsal and 16 ventral segments from 24 newts represented in each sample.

#### Enucleation

During each of the timepoints, animals were anesthetized, and whole eyes were removed using scissors and forceps. Eyelids were left attached at the dorsal side of the eye to serve as marks for proper orientation during embedding and sectioning. Whole eyes were fixed in 4% paraformaldehyde 0.7X PBS overnight at 4°C before processing for histology.

### Single Cell RNA Sequencing Procedures and Bioinformatic Analysis

#### Enzymatic Dissociation of Iris Tissue

PBS was removed from the tubes and replaced with 0.25% trypsin 0.7X PBS without Mg and Ca and with RNAse Inhibitor. Tubes were placed on a Rotator at 37°C for 35 minutes. A P1000 LoBind pipette tip was used to occasionally mix the tissues. The solution was mixed before pipetted through a 70 µm filter and into a BSA-blocked 5mL tube. 0.7X L-15 media containing 5% Fetal Bovine Serum was then added to rinse the filter and deactivate the trypsin. This protocol was optimized to generate at least 80% of alive dissociated single cells assessed using 0.2% Trypan Blue staining.

#### Fixation

Fixation was completed following the Parse Biosciences Cell Fixation Protocol version 2. Dead/alive count was performed after step #6, and final count was performed before adding DMSO in step #16. Following fixation, tubes were placed in a Thermo Scientific Mr. Frosty freezing container at room temperature before stored at -80°C.

#### Sequencing and Bioinformatic Analysis

Single-cell transcriptomic profiling was performed using the Parse Biosciences Evercode™ Whole Transcriptome (WT) kit following the manufacturer’s protocol. Fixed single cells were processed through four rounds of combinatorial barcoding to generate sublibrary pools. Libraries were quantified using Qubit (dsDNA HS Assay) and assessed on an Agilent Bioanalyzer prior to sequencing. Pooled libraries were sequenced on an Illumina NovaSeq 6000 instrument at the Hubbard Center for Genome Studies with the recommended cycle configuration for Parse WT v2 chemistry (Read 1 = 136 cycles for transcript inserts, Read 2 = 86 cycles for barcodes and UMIs, 8 bp i7 and 8 bp i5 index reads) including a 5 % PhiX spike-in. Raw basecall (BCL) files were demultiplexed using bcl-convert v4.2.4. A total of ∼8.8 billion reads were produced across three sequencing runs with an overall sequencing saturation of 0.727. Sequencing depth across the eight sublibraries ranged from 791 million to 1.3 billion reads per sublibrary. The *Pleurodeles waltl* genome assembly and annotations (GCA_026652325, ASM2665232v1) were obtained from the NCBI^13^. The associated GTF annotations were reformatted to ensure a gene_biotype field was present, as required by the Parse BioSciences split-pipe workflow. Annotations were added to previously unannotated genes using BLASTx against the reviewed human proteome (Uniprot ID UP000005640, downloaded August 19, 2024), using an E-value cutoff of 1×10^−4 60,61^. A genome index was created using the Parse BioSciences split-pipe v1.3.1 ‘mkref’ command. Because of the large genome size, the STAR memory limit parameter (lggram) in the mkref.py script was increased to 240 GB. Primary processing and alignment to the genome was completed using Parse BioSciences split-pipe v1.3.1 in ‘mode all’, specifying the WT v2 chemistry and custom newt reference build. split-pipe extracts the combinatorial barcodes and unique molecular identifiers (UMIs) from Read 2, corrects barcodes against the Parse BioSciences whitelist, trims template-switch oligos, removes PCR duplicates, and aligns reads using STAR. After each sublibrary was processed, the outputs were merged with split-pipe ‘comb’. This produced a combined digital gene expression (DGE) matrix, summary reports, and per-cluster differential expression results. Unfiltered count matrices were uploaded to Parse BioSciences’ Trailmaker^TM^ for downstream clustering, visualization, and differential expression analysis.

#### Cell Clustering

Metadata for timepoints (T0, T1, T2, T4, T6, T8, T10, T12, T14, T16) were added manually to each sample. Quality control filters within Trailmaker^TM^ included a cell size distribution filter that selects a minimum number of transcripts per cell, an outlier filter, and a doublet filter set at default settings. Data integration was completed with Seurat as analysis tool^62^, Harmony^63^ as the method, 2000 highly variable genes, logNormalize for normalization, and with 27 number of principal components. Embedding was done using UMAP^64^ with 0.3 minimum distance and cosine distance metric. Clustering was done using Leiden with 0.8 resolution. Mitochondrial content analysis was performed by aligning reads using Parse BioSciences split-pipe against the reference *Pleurodeles waltl* mitochondrial DNA. Table S1 contains relevant statistics and parameters for the samples. Marker genes were selected based on the genes enriched in each cluster and existing literature for known cell types. “Custom cell sets” were generated by merging Leiden clusters into distinct cell types with known and distinct marker genes (see Figure 1 D-E). For the different analysis used, cells were reclustered into three new subsets: “Day 0 IPE, NPCE, and PCE Only”, “Dorsal IPE Only”, and “Macrophages Only”. “Day 0 IPE, NPCE, and PCE Only” cells were the intersection of T0 cells and the IPE, NPCE, and PCE cells from “custom cell sets”. “Dorsal IPE Only” cells were the intersection of the IPE “custom cell set” and VAX2 expression < 1 to remove the majority of ventral IPE cells. VAX2 was used rather than TBX5 due to reprogrammed LECs losing TBX5 expression (Figure S2B and Table S5). “Macrophages only” cells were derived from “custom cell sets”. “Engulfing_Macrophages_CSF1R+” cells were derived by the intersection of Leiden clusters 9 and 5, and CSF1R expression > 1 from the “Macrophages only” subset. Reclustering was performed in Trailmaker^TM^ with default parameters.

#### Analysis using Trailmaker^TM^

Trailmaker^TM^ was used for all categorical and continuous UMAPs, dot plots, volcano plots, and frequency plots (Figure 1D-F, Figure 3B-D, Figure 4A-E, Figure 6 C-D, F-G). Dot plots used in Figure 1E and 3C were generated using custom gene markers. Dot plots in Figures 4C and 6D were generated using the marker gene feature per cluster. Volcano plots had a significance cutoff of adj p-value < 0.05, and expression cutoff of logFC < -0.6 and logFC > 0.6. Batch Differential Expression tables were derived from Trailmaker^TM^ (Tables S2, S3, S5, S7). Data were exported as Seurat objects for downstream analysis in R studio.

#### Pearson Correlation, Cell Cycle, and PCA Analysis

Pearson correlation heatmaps were generated using the Pheatmap package (Figure 2A-C, Figure S2A). Cell cycle analysis visualized as kernel density plots (Figure S2C) and as stacked bar graphs (Figure 2E and Figure 6J) were generated using the Tricycle and ggplot2 packages^65^. PCA analysis for Figure 2C was generated using Seurat and visualized using ggplot2. Dorsal and ventral IPE cells were subsetted by timepoint before being both normalized and scaled.

#### Enrichr Transcription Factor Prediction

Transcriptional factors were predicted using the Transcription Factor PPIs database on Enrichr^25^. For Figure 2F the input list was derived from the Batch Differential Expression records of each cell type at each stage (Table S3) with cutoffs logFC >0.6 and adj p-value <0.05. For Figure S5E, the input list was composed of all the genes listed in each gene module from Figure S5D and Table S9.

#### Venn Diagrams

Venn diagrams were generated using Venny 2.0^66^. In Figure 3 the input lists for each Venn diagram were derived from the volcano plot comparisons (Table S6) with cutoffs adj p-value <0.05 and logFC <-0.6 or >0.6 for under- or over-represented genes, respectively.

#### R-squared Value Analysis

For each cluster in the “Dorsal IPE Only” cell set, the Batch Differential Expression datasheets were filtered for genes expressed in more than 20% of cells in that cluster (pct_1 > 20). R-squared value was then computed based on the expression of all filtered genes in each cell. For Figure 4H, the top 4 positively and negatively correlated pairs were visualized using ggplot2 in R.

#### Gene Ontology Enrichment Analysis

Genes were filtered for adj p-value < 0.05 and logFC >0.6 from Batch Differential Expression datasheets derived from “custom cell sets” per stage (Table S3) or individual clusters (Table S5). GO enrichment analysis was performed using DAVID with the background set to all available annotated newt genes^67^. Tables S4 and S8 contain the full list of enriched GO terms for the two analyses performed. A comprehensive selection of enriched GO terms was plotted using ggplot2 for Figure 2D and Figure 4G.

#### Pseudotime Trajectory Analysis

The “Dorsal IPE Only” subset was loaded to R studio as a Seurat object and the Monocle3 package^68^ was used to generate the initial three partitions (Figure S5A). The two smaller partitions which mainly included T0 cells were excluded to allow the trajectory analysis. A subset of the bigger partition, which included the remaining timepoints, was used to run the trajectory analysis. The subset was made by choosing cells from T1 to the PROX1+ LECs as shown in Figure S5A, B and C. A categorical plot with timepoints (Figure 5A), a continuous plot with PROX1 expression (Figure 5B), and a pseudotime plot with node labels (Figure 5C) were all generated from this new subset. A heatmap of the top three marker genes per node was generated across pseudotime by taking the average pseudotime value of the cells associated with a node and placing them in a pseudo-chronological order using Monocle3, Pheatmap, and Matrix packages (Figure 5D). Gene modules were then generated using Monocle3 (Figure S5D). Linear expression vs. pseudotime graphs were plotted using Monocle3 (Figure 5F).

#### CellChat

The CellChat package^69^ was run in R studio using the Seurat object containing all cells from the project. The chord diagram in Figure 6A that depicts all cell-to-cell interactions was generated using CellChat. The bubbleplot in Figure 6B was made using ggplot2 with selected interactions extracted from the CellChat object.

*Histology*

#### Histological Timecourse Preparation and Imaging

Paraformaldehyde-fixed timepoint-matched whole eyes from sibling animals were dehydrated in an alcohol gradient up to 100% followed by xylene and paraffin washes. Eyes were then embedded individually in molds for sectioning in the sagittal plane. 5µm sections were cut using a rotary microtome. Three slides per timepoint were stained using hematoxylin and eosin (H&E) using standard protocols. Images were taken using a Nikon A1R HD confocal microscope at high magnification. High-magnification, high resolution images in Figure 1A were stitched using the Nikon’s NIS Elements software.

## Figures and Figure Captions

**Figure S1: UMAPs for samples and marker genes**

A) UMAPs for custom cell sets per timepoint

B) UMAPs for custom cell set marker genes

C) UMAP for PAX6 expression

D) GLM analysis of cell type abundance over time

**Figure S2: Correlation heatmaps and cell cycle analysis**

A) Correlation heatmaps across timepoints for all cell types

B) UMAPs for TBX5 and VAX2 expression for IPE identification

C) Cell cycle analysis using tricycle for dorsal and ventral IPE per timepoint

**Figure S3: Gene expression in PAX6+ intact iris cells**

A) UMAPs for IPE, NPCE, and PCE subpopulation marker genes

**Figure S4: Gene expression and IPE subpopulation abundance across timepoints**

A) Heatmap with top genes per timepoint for dorsal IPE cells

B) GLM analysis of dorsal IPE clusters over time

**Figure S5: Gene expression patterns over the pseudotime trajectory**

A) Monocle3 generated UMAP pseudotime trajectory with PROX1 expression

B) Monocle3 generated UMAP pseudotime trajectory with actual timepoints

C) Selected trajectory path with high correlation to timepoints sampled

D) Gene modules with different gene expression patterns along the trajectory

E) Predicted transcriptional factors regulating genes in each module from D

**Figure S6: Cell-to-cell interactions and macrophage analysis**

A) Selected cell-to-cell interactions derived from CellChat

B) Frequency plot for macrophage clusters over timepoints

C) Secondary doublet analysis for macrophage clusters

## Tables and Table Captions

**Table S1: Quality Metrics & Project Details**

**Table S2: Batch Dicerential Expression Data for Custom Cell Sets**

**Table S3: Batch Dicerential Expression Data per Custom Cell Set by Regeneration Stage**

**Table S4: GO Term Enrichment Analysis for Custom Cell Sets by Regeneration Stage**

**Table S5: Batch Dicerential Expression Data for “Day 0 IPE, NPCE, PCE Only” Subset**

**Table S6: Gene Expression Comparisons for Selected Intact Clusters**

**Table S7: Batch Dicerential Expression Data for “Dorsal IPE Only” Subset**

**Table S8: GO Term Enrichment Analysis for Dorsal IPE Clusters**

**Table S9: Full Gene List for Gene Modules from Monocle3 trajectory**

**Table S10: Batch Dicerential Expression Data for “Macrophages Only” Subset**

