## Supplementary figures and images for "A Single Cell Atlas of the Newt Iris During Lens Regeneration"

### Figure S1

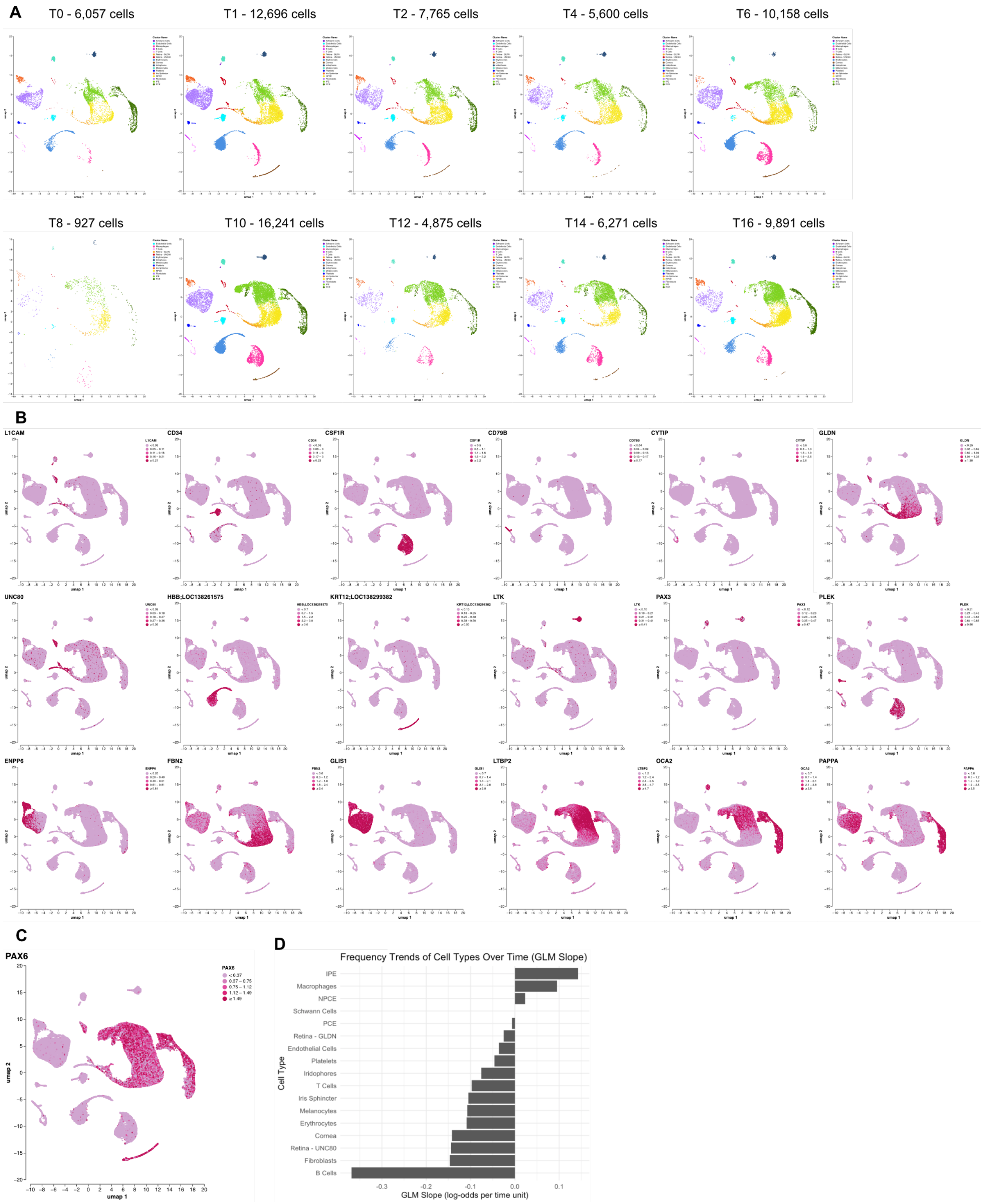

### Figure S2

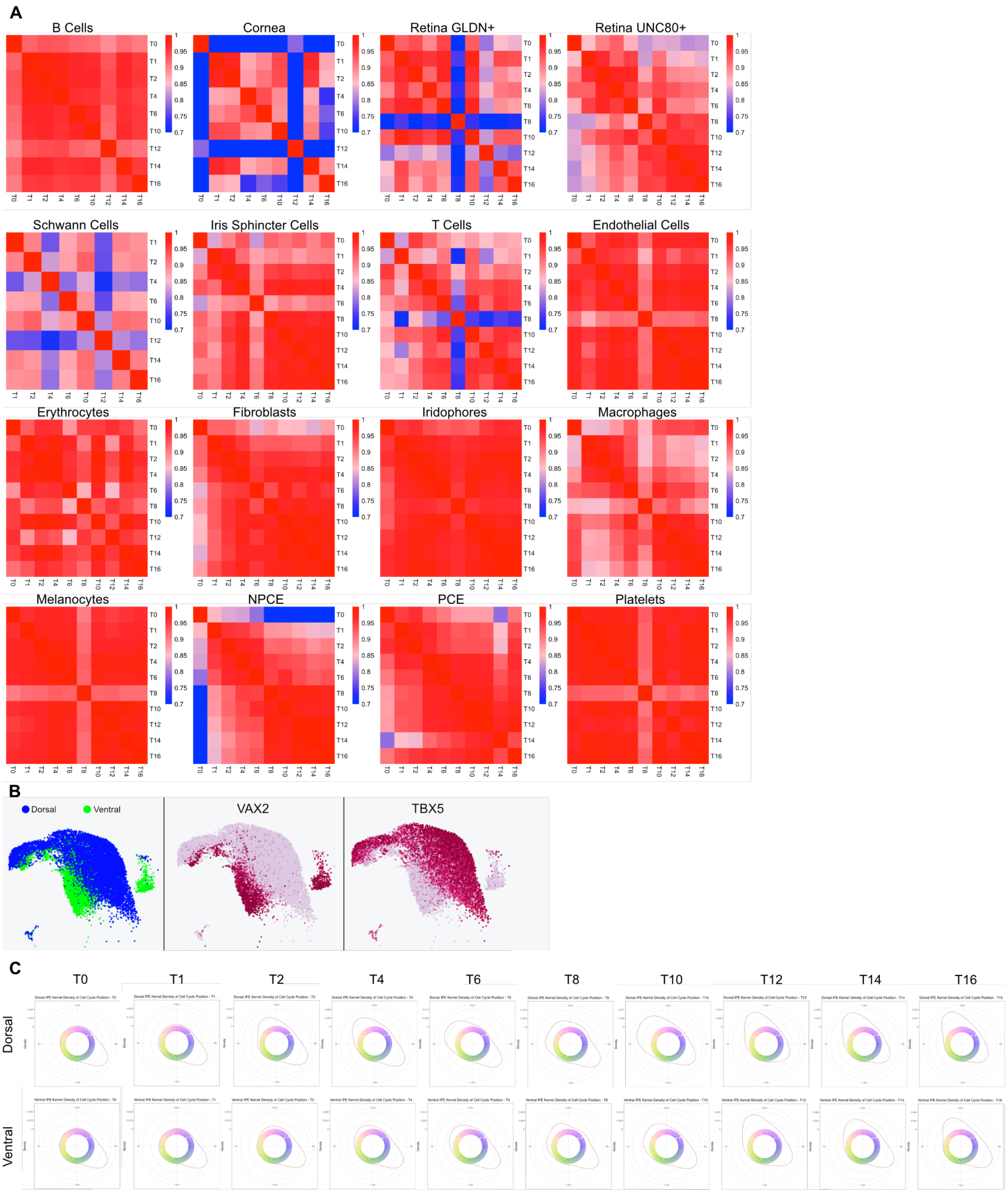

### Figure S3

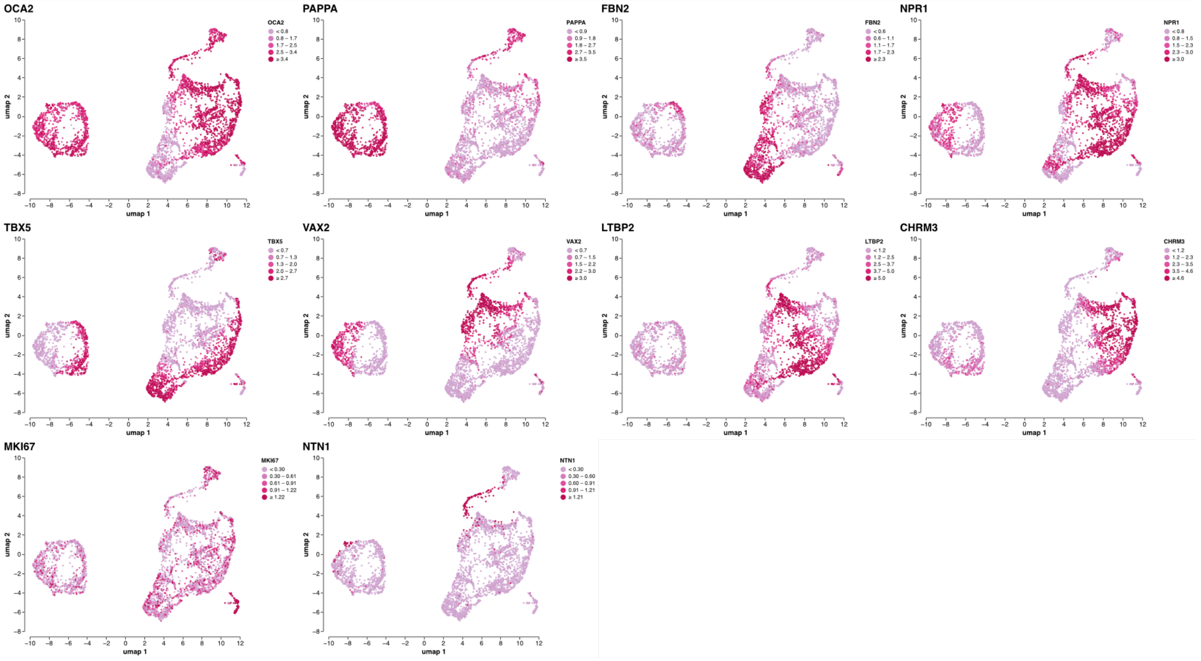

### Figure S4

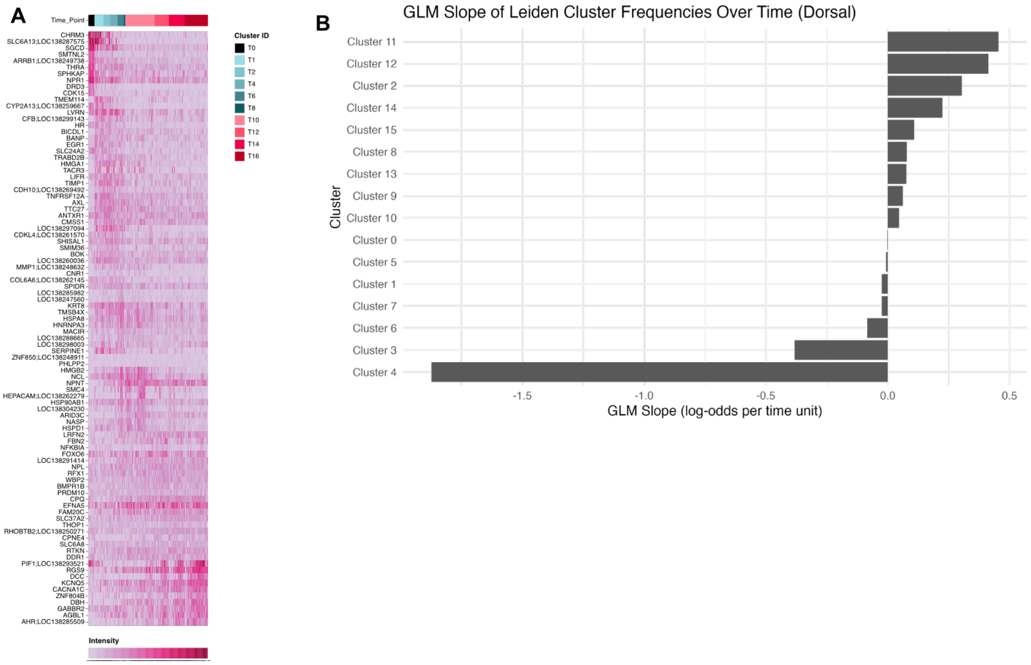

### Figure S5

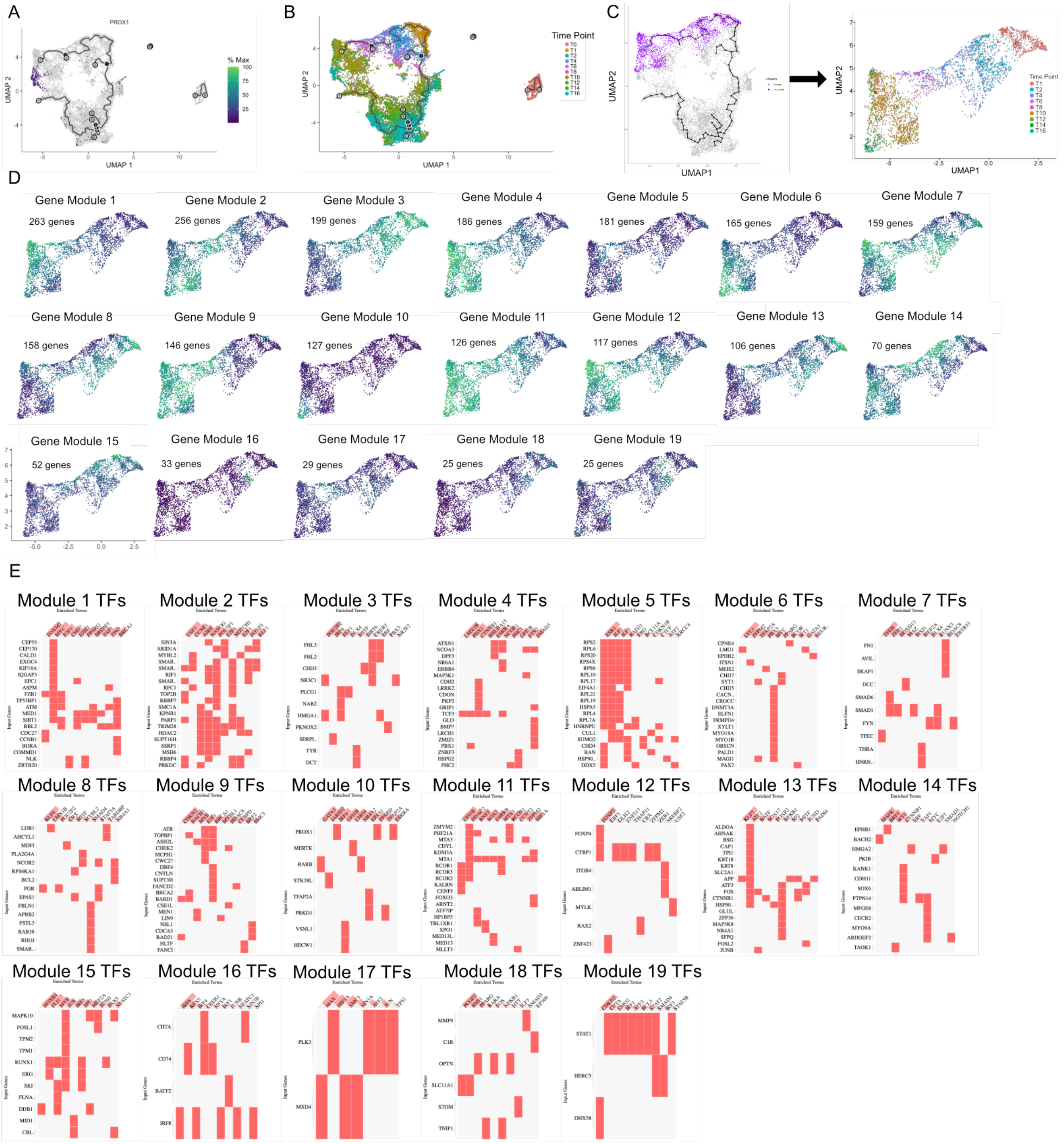

### Figure S6

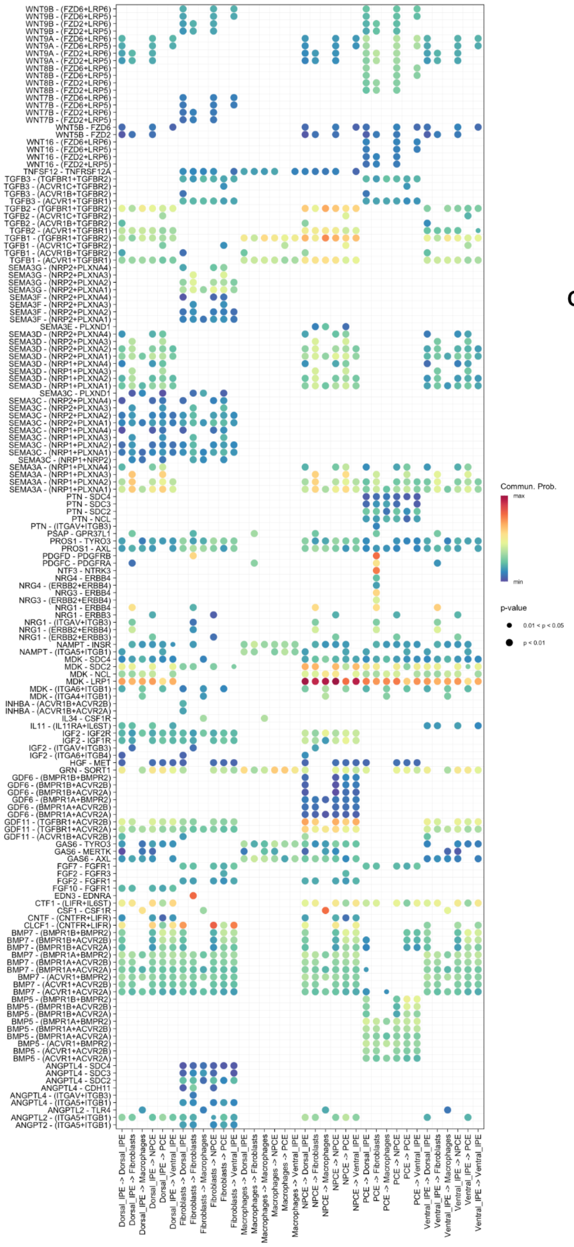

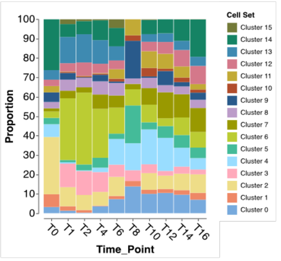

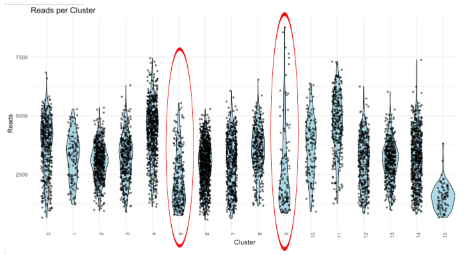
